# AmpliCI: A High-resolution Model-Based Approach for Denoising Illumina Amplicon Data

**DOI:** 10.1101/2020.02.23.961227

**Authors:** Xiyu Peng, Karin Dorman

## Abstract

**Motivation:** Next-generation amplicon sequencing is a powerful tool for investigating microbial communities. One main challenge is to distinguish true biological variants from errors caused by PCR and sequencing. In the traditional analysis pipeline, such errors are eliminated by clustering reads within a sequence similarity threshold, usually 97%, and constructing operational taxonomic units, but the arbitrary threshold leads to low resolution and high false positive rates. Recently developed “denoising” methods have proven able to resolve single-nucleotide amplicon variants, but they still miss low frequency sequences, especially those near abundant variants, because they ignore the sequencing quality information.

**Results:** We introduce AmpliCI, a reference-free, model-based method for rapidly resolving the number, abundance and identity of error-free sequences in massive Illumina amplicon datasets. AmpliCI takes into account quality information and allows the data, not an arbitrary threshold or an external database, to drive conclusions. AmpliCI estimates a finite mixture model, using a greedy strategy to gradually select error-free sequences and approximately maximize the likelihood. We show that AmpliCI is superior to three popular denoising methods, with acceptable computation time and memory usage.

**Availability:** Source code available at https://github.com/DormanLab/AmpliCI

## 1. Introduction

High throughput sequencing has revolutionized the study of microbial communities. A common strategy to identify and quantify species present in a sample is to amplify and sequence biomarker genes, like 16S rRNA or fungal Internal Transcribed Spacer (ITS). These biomarker genes contain both conserved and hypervariable regions, allowing amplification while still distinguishing organisms. Amplicon sequencing is a powerful tool for studying the composition and structure of microbial communities.

The utility of biomarkers is degraded by sequencing errors, PCR amplification errors, and intra-strain/species-specific variability [1]. To account for these factors, a typical first step of microbiome analysis is to resolve the data into Operational Taxonomic Units (OTUs), or clusters of sequences with 97% or greater similarity. There are many methods for identifying OTUs [2], roughly classifiable into closed-reference methods, which use a reference database of known organisms, or *de novo* methods. However, when applied to mock communities, it is widely found that both types of methods cannot accurately identify true OTUs in a sample [3, 4, 5, 6, 7, 8].

OTUs are problematic entities, lacking both biological and physical interpretability. They only roughly correspond to biological species, genera or higher taxonomic entities, and they do not correspond to true, error-free sequences in the sample. Thus, OTU-based methods are prone to both false positives and negatives, reporting error sequences as OTUs and missing subtle and real biological sequence variation, such as SNPs. The 97% threshold, motivated by empirical studies [9, 10], fails to reliably achieve genus or species level resolution [11, 12]. There are distinct species with 97% or more similar 16S rRNA [13, 14], and strains whose 16S rRNA locally differ by more than 3% [15].

Amplicon sequencing data from current Illumina platforms support *de novo* single-nucleotide resolution [16]. Modern methods attempt to identify all the unique sequences in the sample [17, 18, 19, 20, 16, 21, 22, 23]. Such denoising methods make no biological judgment on taxonomic entities, but simply remove or correct sequences produced by sequencing errors or, sometimes, PCR errors. The denoised sequences are called Amplicon Sequence Variants (ASVs) [16], sub-OTUs (sOTUs) [22], or zero-radius OTUs (ZOTUs) [21]. Their higher resolving power, lower false positive rates, and greater inter-sample consistency have made denoising methods the recommended tool for biomarker gene analysis [2, 1, 8].

There are currently three widely used denoising methods [8]: DADA2 [16], UNOISE3 [21] and Deblur [22]. UNOISE3 and Deblur ignore the quality information and greedily select true sequences assuming conservative error rates. DADA2 uses a greedy, hierarchical divisive clustering algorithm based on a probabilistic error model, while accounting for averaged quality score information. Only DADA2 infers error rates from data, a potential advantage, since sequencing machines and PCR conditions affect error profiles [18].

We introduce AmpliCI, Amplicon Clustering Inference, a model-based algorithm for denoising Illumina amplicon data. The statistical model underlying AmpliCI is a finite mixture model, which for computational feasibility, is maximized using an approximate, greedy scheme. Like DADA2, AmpliCI uses a formal model for sequencing errors, but it retains higher resolving power by not averaging quality scores among reads with identical sequences. AmpliCI considers both substitution and indel errors, estimating error parameters directly from the sample. We test our method on simulated, mock and real datasets. AmpliCI beats current algorithms, particularly achieving higher accuracy for highly-related sequences.

## 2. Method

### 2.1. Statistical Model

We start with read set 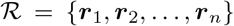 and quality set 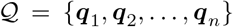, containing *n* sequences of base calls and accompanying quality scores. We assume the data are independent draws from a *K*-component mixture distribution, where the *k*th component is generated by the true sequence (haplotype) ***h***_*k*_. The likelihood function for fixed *K* is

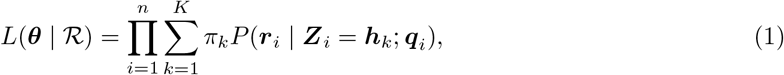

where ***Z***_*i*_ is the unknown source sequence of read ***r***_*i*_, parameters 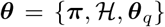 are the mixing pro-portions ***π*** = (*π*_1_, *π*_2_, …, *π*_*K*_), the set of true haplotypes 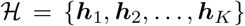, and parameters ***θ***_*q*_, dictating how the quality scores ***q***_*i*_, treated as observed covariates, impact the likelihood of read ***r***_*i*_.

The conditional probability *P*(***r***_*i*_ | ***Z***_*i*_ = ***h***_*k*_; ***q***_*i*_) or transition probability for generating read ***r***_*i*_ from haplotype ***h***_*k*_ in the context of the quality scores ***q***_*i*_, is calculated conditional on the pairwise alignment between the read ***r***_*i*_ and the source haplotype ***h***_*k*_. Since substitution errors strongly exceed insertion or deletion (indel) errors in Illumina sequencing [24], we use a simple model to penalize indels. We assume an insertion can occur *before* or a deletion *at* any one of the *l*_*k*_ positions in the *k*th haplotype at a very small, constant rate *δ*. Assuming these events are independent, the number *d*_*i*_ of observed indel events in the *i*th read may be approximately modeled as a truncated Poisson distribution,

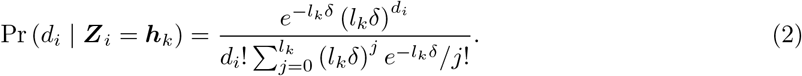

We ignore indel lengths, assuming all plausible lengths are equally likely. Technically, because reads have fixed lengths, indels are neither independent nor their lengths ignorable, but the approximation should be good for short indels, small *δ*, and by treating indels after the last haplotype position as necessary consequences of earlier indels. Finally, assuming errors are independent across sites, the transition probability is

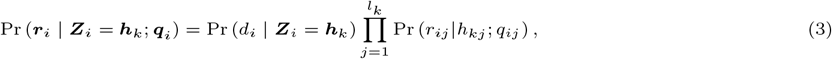

where *j* indexes positions in haplotype ***h***_*k*_ and the *aligned* read/quality sequences. The term Pr(*r*_*ij*_ |*h*_*kj*_; *q*_*ij*_) is the probability of generating nucleotide *r*_*ij*_ from *h*_*kj*_ with quality score *q*_*ij*_ at alignment position *j*, understood to be 1 when *r*_*ij*_ is a deletion. Like DADA2 [16], we estimate the log probabilities log Pr(*r*_*ij*_ |*h*_*kj*_; *q*_*ij*_), for each choice of *h*_*kj*_ and *r*_*ij*_ in {A, C, G, T}, using LOESS regression on the quality score (details in §2.4).

### 2.2. Greedy Haplotype Selection

Maximizing likelihood (1) is extremely challenging [25]. Instead, we make two key assumptions that motivate an approximate maximization scheme. Capitalizing on the low error rates of Illumina sequencing, our assumptions are: (1) all true haplotypes are observed at least once without error and (2) errored reads that match a true haplotype are overwhelmingly sourced from more abundant haplotypes. Under the first assumption, if we could know the true proportion *η*_***s***_ = Pr(***Z***_*i*_ = ***s***) for all unique sequences *s* in the sample, then the sequences with positive proportion are true haplotypes. We use the other assumption to find approximate, rapid estimates of *η*_***s***_ for all unique ***s***.

Suppose the true set of haplotypes is 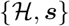 for some unique sequence 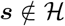. The random observed abundance *A*_***s**o*_ of ***s*** is comprised of the true abundance *A*_***s**t*_, plus the number 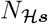 of misreads of other haplotypes as read sequence ***s***, and minus the number 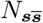 of misreads of sequence ***s*** (Supplementary Figure S1),

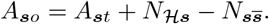

If we take the expectation of both sides, then

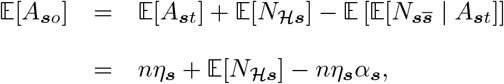

where *α*_***s***_ is the probability of misreading sequence ***s*** with at least one error. If we assume the misread probability is a constant *α* independent of the source sequence ***s***, then a method-of-moments estimator of the expected true proportion is

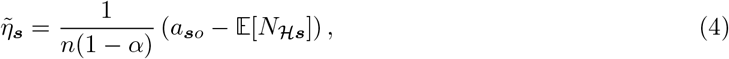

where *a*_***s**o*_ is the observed abundance of sequence ***s***. We could use (4) to estimate *η*_***s***_ for all unique 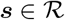, but the equation relies on knowing the haplotypes. We use our key assumptions along with ideas from Deblur [22] and UNOISE2 [21] to incrementally find haplotypes.

For convenience, we reindex the whole data set. Since unique sequences with very low abundance are unlikely to be haplotypes and are certainly difficult to distinguish from low-level contaminants, we remove all unique sequences with abundance below a threshold *c* from consideration (*c* also appears later in contamination detection). We are left with *M* unique sequences 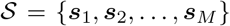, sorted from highest to lowest observed abundances 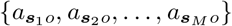. The clustering of reads by unique sequence induces a partition on the read set 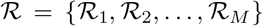 and quality set 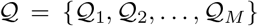. Subset 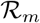 contains reindexed reads ***r***_*mi*_ = ***s***_*m*_, and subset 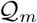 contains the accompanying reindexed quality score sequences ***q***_*mi*_, for 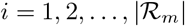. DADA2 similarly rearranges the dataset, but averages quality scores within the *m*th subset [16]. Retention of original quality scores is an important distinction, increasing sensitivity by allowing AmpliCI to detect members of 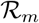 that are likely misreads of other haplotypes.

Let 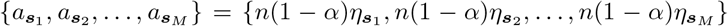 be the expected scaled true abundances (shortened to scaled abundances) we aim to estimate. Assuming the most observed unique sequence is a haplotype and no other haplotype is misread as this unique sequence, we start by setting 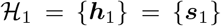 and method of moments estimator 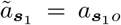. Given *k* haplotypes so far, the (*k* + 1)th haplotype will be chosen from 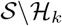. We consider each 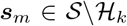 as a possible haplotype and compute the scaled abundance estimate

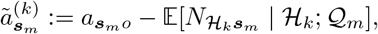

where we condition on the current haplotype set 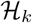 and utilize the observed quality scores 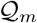 of the reads matching the *m*th unique sequence ***s***_*m*_. This estimate is obtained under the assumption that the haplotype set is 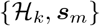, which is clearly incorrect, especially in early iterations. However, the approximation is essential for computation, as witnessed by attempts to estimate a complete model in a simpler context [26], and it is reasonable under our second key assumption. Further derivations in Supplementary Material S1 yield estimation function

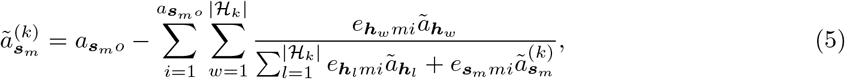

with *e*_***s**mi*_ = Pr(***s***_*m*_ | ***Z***_*mi*_ = ***s***; ***q***_*mi*_) given by (3). Estimate 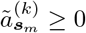 is a fixed point of (5) (Supplementary Material S2), which can be found through fixed point iteration (Supplementary Material S3).

We now describe the algorithm for approximately maximizing (1) (Fig. 2). The estimated scaled abundance 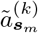 at iteration *k* estimates the expected number of error-free reads of sequence ***s***_*m*_ and is a suitable statistic for selecting the next candidate haplotype. The sequence with highest scaled abundance is not only a likely haplotype, but also the haplotype most likely distorting the observed abundance of other candidate haplotypes. Therefore, at each iteration *k*, we propose candidate ***h***_*k*+1_ as the sequence with highest 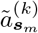, but whether we accept the proposed candidate is a model selection issue.

**Figure 1:**
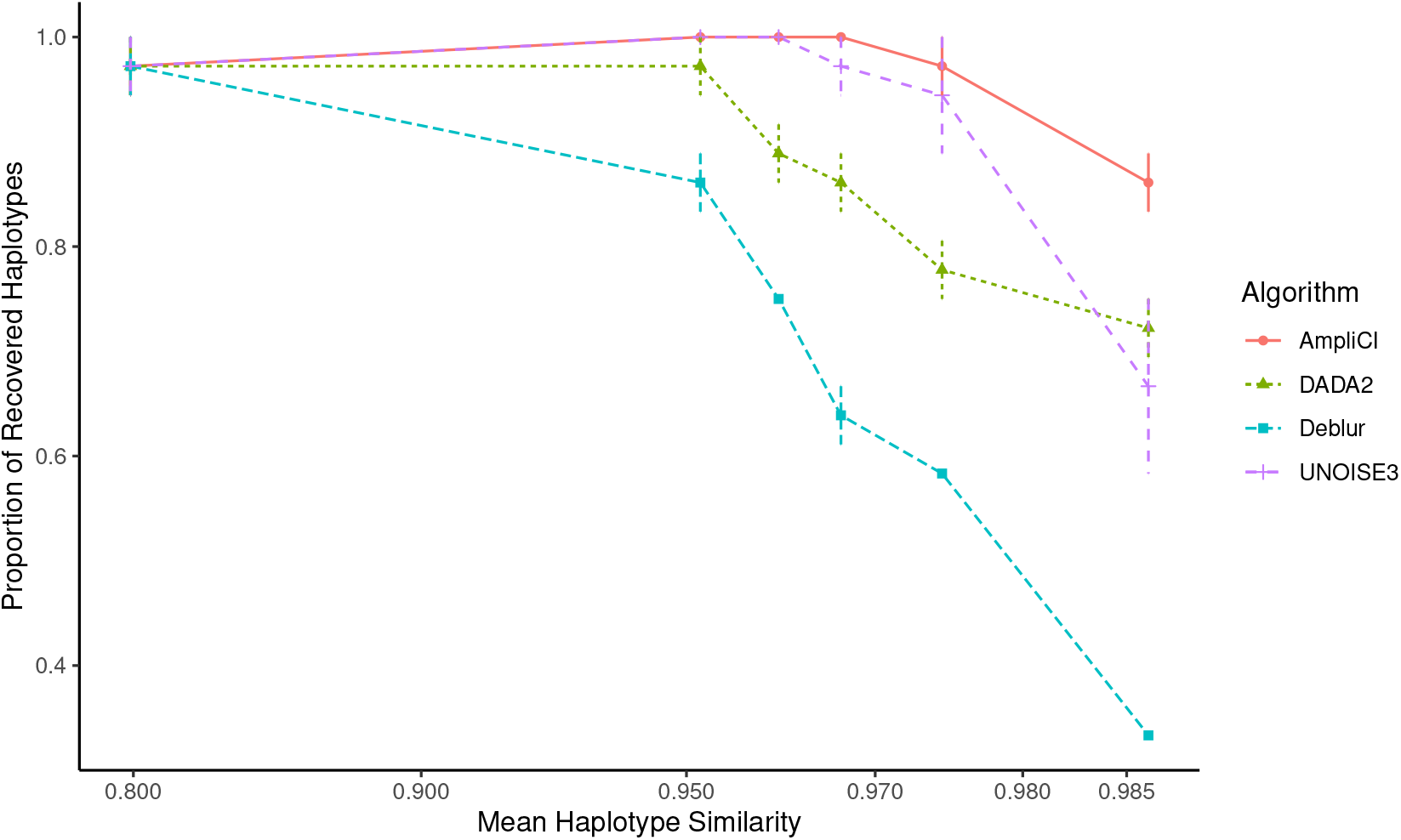
Mean proportion of detected haplotypes ± standard error for six levels of simulated haplotype similarity. For each level, the *x* coordinate is set to the mean haplotype similarity of the three simulated datasets. When the similarity level among haplotypes increases, AmpliCI performs better than competing methods. Detailed results on each dataset are provided in Supplementary Table S1.

**Figure 2:**
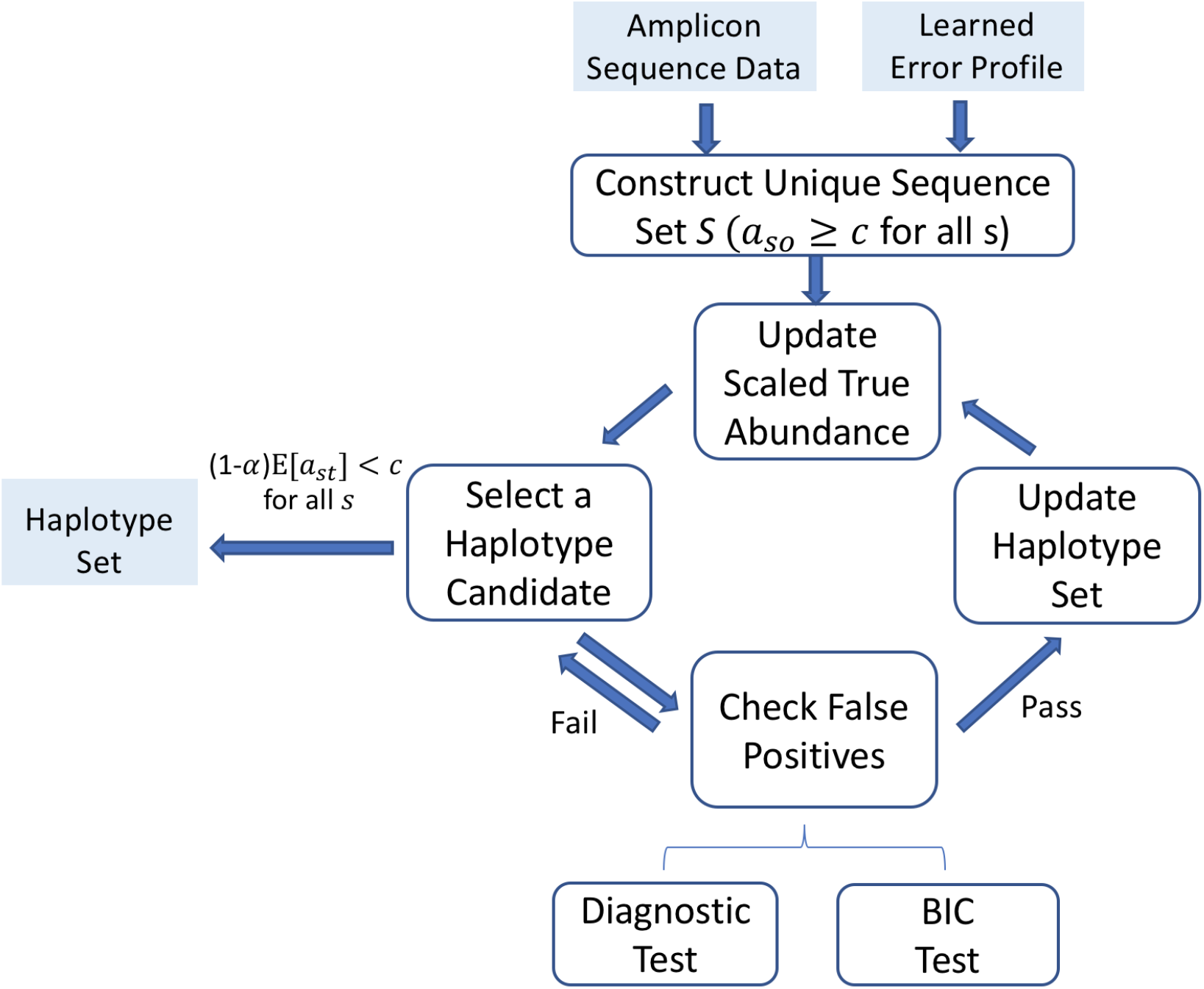
AmpliCI: inferring ASVs from samples. 1) A hash index is applied to construct the unique sequence set S, and the haplotype set 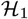 contains the most abundant unique sequence. 2) Given the current haplotype set 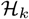, the scaled abundances 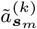 are estimated for each remaining unique sequence ***s***_*m*_ via update function (5), and the haplotype candidate 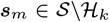 with the highest scaled abundance is selected. 3) Test if the diagnostic probability is smaller than the given threshold and the approximate Bayesian Information Criterion (BIC) improves. If the haplotype is discarded, we select the next most abundant candidate. Otherwise, add candidate ***s***_*m*_ to the current haplotype set, 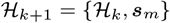. 4) repeat 2) - 3) until the scaled abundance 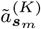 of all remaining unique sequences 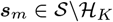 are below the user-determined lower bound *c*. 5) Output the haplotype set 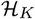 with *K* haplotypes.

To assess model goodness of fit with ***s***_*m*_ in the haplotype set, we compute an approximate Bayesian Information Criterion (BIC) (Supplementary Material S5) and permanently reject candidate haplotypes that do not improve the BIC. Surviving candidates are then diagnosed for contamination (Supplementary Material S6). Briefly, we assume contamination introduces *z* (default *z* = *c* − 1) copies of the candidate ***s***_*m*_. If it is “easy” to generate all 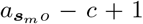 remaining copies as misreads of haplotypes in 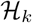, then we doubt whether ***s***_*m*_ is a haplotype. To quantify the ease of this event, we compute 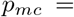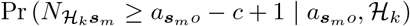 using the fact that the misreads 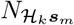 follow a Poisson Binomial distribution. Small values of the probability indicate it is *not* easy to explain the observed count without haplotype ***s***_*m*_. By default, we require *p*_*mc*_ < 0.001/*M* for *M* unique candidate sequences.

When a candidate fails to improve the BIC or its diagnostic probability exceeds the threshold, we repeatedly consider next most abundant candidates until a haplotype is accepted or no candidate remains. If a new haplotype is accepted, the haplotype set 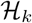 is updated to 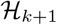, and the process repeats. The algorithm continues iterating until there are *K* haplotypes in 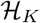 and all scaled abundances 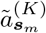 of remaining candidate sequences fall below the threshold *c*.

### 2.3. Abundance Estimation

An important secondary goal of a denoising algorithm is abundance estimation, which is required for chimaera detection [27] and many downstream analyses [1]. AmpliCI simply uses the final, estimated scaled abundances 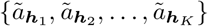 for this purpose. Under the assumption of sequence-independent misread rate *α*_*s*_ = *α* for all 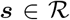, these values are directly proportional to the true haplotype abundances. However, with a fully probabilistic model such as AmpliCI, it is possible to imagine more sophisticated methods for abundance quantification. These and related issues are discussed in Supplementary Material S7.

### 2.4. Implementation

We implement AmpliCI in the C language. By default, the insertion rate is 4 × 10^−5^ and deletion rate 2 × 10^−5^, consistent with previous estimates [24]. Here we describe the estimation of the remaining error parameters ***θ***_*q*_ and how we avoid the computational expense of all-against-all alignments.

It is well known that quality scores do not perfectly predict error rates [18], but they are strongly correlated with the presence of errors [16], even if the exact relationship can vary by dataset [28]. In our experience, it is very important to estimate the error properties of each sample before denoising. AmpliCI independently learns the error profile for each sample, after demultiplexing (Supplementary Material S4). Briefly, error rates are initialized to those dictated by Phred quality scores [29], *i.e.*,

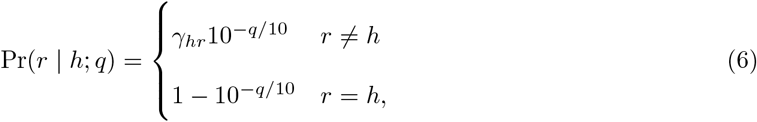

where 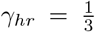 for all *h, r* ∈ {A, C, G, T}, *h* ≠ *r*. After using AmpliCI logic to add a sequence to the haplotype set, new error rates are estimated by weighted LOESS regression assuming the current haplotype set. We continue to add haplotypes and re-estimate error rates until they stabilize, then use the final estimates in a second run of AmpliCI without error rate estimation.

Since the computational complexity of the Needleman-Wunsch algorithm for alignment (see §2.1) is high, we implement an *alignment-free* strategy (by default), where transition probabilities between reads and haplotypes are calculated without pairwise alignment. The *alignment-free* strategy decreases the average runtime on simulated datasets (§2.5) from 0.89s (`-z` option) to 0.06s. It works because sequencing indel errors are rare, but when they occur, the alignment-free approach will select indel-containing reads when threshold *c* is low. To mitigate this problem, once a haplotype candidate is selected, we recalculate its scaled abundance based on the pairwise alignments (scores: gap = −5, match = 2, mismatch = −2 for transition, −3 for transversion) to each current haplotype. If the scaled abundance drops below the threshold *c*, which is expected for indel misreads, the candidate haplotype is permanently dropped.

### 2.5. Data Simulation

Synthetic datasets are simulated from a model related to that of read generator ART [30]. For the simulation with most easily detected haplotypes, we simply use the *K* = 12 most abundant haplotypes from the Extreme mock dataset [16], but for the other synthetic datasets, we first simulate 12 haplotypes on a star-shaped phylogeny, under the Jukes-Cantor model [31] initialized with the consensus sequence of the 12 most abundant Extreme sequences. The similarity between haplotypes is controlled by the branch length *c*_*d*_ in the star phylogeny (Supplementary Table S1), plus the requirement that all haplotypes are distinct. The relative abundance of the 12 haplotypes are (0.377, 0.278, 0.101, 0.052, 0.049, 0.04, 0.039, 0.036, 0.006, 0.005, 0.005, 0.004), implying unbalanced clusters. Then, for each read, we randomly pick a haplotype among the 12 with the given mixing proportions, simulate the quality score sequence based on a read position-specific quality score empirical distribution, and given the haplotype, simulate nucleotide sequences independently assuming Phred quality scores (6). The mixing proportions, quality score distribution and *γ*_*hr*_ in the Phred error probabilities are the maximum likelihood estimates obtained by EM optimization of the mixture model (1) for 3,000 random reads from the extreme mock dataset [16] with fixed, known haplotypes and initialized with *γ*_*hr*_ = 1/3. The insertion and deletion rates are set to 2 × 10^−5^ per position. The simulation error model is related to but not identical to the AmpliCI error model, and importantly, we do not enforce the additional AmpliCI assumptions. We simulate 3,000 reads per synthetic dataset at five different similarity levels.

### 2.6. Analysis of Mock and Real Datasets

We examine three mock datasets, Extreme [16], Even1 and Stag1 from Mock5 [32, 33], and one real dataset of vaginal microbiota [34]. A pertinent summary of all datasets is given in Table 2. Most denoisers are part of a complete analysis pipeline, including both pre- and post-processing steps. The pipeline we use is in Figure S2. To facilitate comparison, especially for mock data, we equalize as much as possible in these pipelines, though see Supplementary Material S9. Deviations in the analysis for the real dataset are described in the results.

Only forward reads of each sample are considered as input to the denoisers. Even1 and Stag1 are demultiplexed out of the Mock5 dataset using QIIME1 [35] `script_split libraries_fastq.py` and default settings. For the real dataset, we download the already demultiplexed data. All reads are truncated at 240nt and shorter reads are discarded. We also remove reads that contain any quality score less than 3 or ambiguous nucleotide ‘N’. The resulting datasets are used as the *same* input for all four algorithms to test their performance. However, Deblur [22] includes a positive pre-filtering step to remove reads with no match in reference database 88% OTUs from Greengenes v13.8 [36], by SortMeRNA [37]. Since we could not turn off this feature, we implement a version of AmpliCI with positive *post*-filtering against the same database.

Denoiser output is often processed to remove chimaera sequences and other artifacts. For mock datasets, we use the UCHIME3 *de novo* method [27] to remove chimaeras for all methods except UNOISE3, which uses an embedded version of UCHIME3 *de novo*. A conservative version of AmpliCI, called AmpliCI-con, uses post hoc filtering of detected haplotypes after chimaera removal, discarding haplotypes whose diagnostic probabilities *p*_*mc*_ for contamination exceed 10^−40^.

## 3. Results

We compare AmpliCI with three prevalent denoising methods: DADA2 [16], UNOISE3 [21] and Deblur [22]. Summary information, including version number, of all algorithms is given in Supplementary Table S3.

### 3.1. Simulated Datasets

We compare the algorithms on synthetic data, using a common abundance threshold *c* = 2, the minimum dictated by DADA2, and all other default parameters. After verifying that Deblur’s pre-filtering step removes no simulated haplotype, we assess the methods’ ability to recover true haplotypes and read assignments. Since Deblur does not have a read assignment method, we use USEARCH v11.0 [38] to assign reads to Deblur haplotypes. For AmpliCI, we assign reads to the cluster with maximum posterior assignment probability (See Eq. (S5) in Supplementary Methods S7). To assess the accuracy of read assignments, we compute the Adjusted Rand Index (ARI) [39].

Figure 1 and Supplementary Table S1 show the performance of all methods on three replicate datasets under six simulation conditions, where we vary the similarity of the haplotypes. For datasets with 0.80 mean haplotype similarity, all algorithms perform well, although UNOISE3 identifies two false haplotypes in one of the datasets. As the mean haplotype similarity increases, performance declines for all algorithms, but for datasets with haplotype similarity greater than 0.965, AmpliCI achieves the best performance among all three datasets, with the highest ARI and number of recovered haplotypes.

### 3.2. Mock Datasets

We analyze the forward reads of three mock datasets, real samples of known microbial communities, which have been widely used for microbiome method benchmarking. To screen low abundance contaminants and handle the peculiar noise patterns of real data, denoising algorithms take extra steps to screen putative haplotypes. All algorithms, including AmpliCI, overestimate the chance of read errors (see Discussion). In addition, Deblur and UNOISE3 recommend setting a high abundance threshold. While a high threshold can reduce the number of false positives, it also reduces method sensitivity. In contrast, DADA2 sets a low threshold of 2 and tests the evidence in support of candidate haplotypes, accepting new haplotypes only if its diagnostic probability falls below 10^−40^, a highly conservative choice. Deblur further reduces false positives by positive prefiltering all reads against an outside database [36]. AmpliCI can use all these strategies to reduce the false positive rate. We use both a low (2) and high (10) abundance threshold on all methods, except DADA2. A conservative version of AmpliCI, AmpliCI-con, sets a conservative threshold on diagnostic probabilities (10^−40^) in the contamination screen. We try positive filtering on AmpliCI results whenever Deblur’s performance is superior.

Table 1 shows the result when true positives (TP) only include estimated haplotypes that are a 100% match to the provided mock reference sequences. AmpliCI achieves more or equal true haplotypes with fewer false haplotypes. With low abundance threshold 2, AmpliCI-con, with conservative threshold, performs better than DADA2 and UNOISE3 on all three datasets. Deblur is better than AmpliCI on the Even1 dataset, but only with the assistance of positive filtering. AmpliCI, though designed for fine-scale resolution, continues to perform best at high abundance threshold 10 on the Extreme and Even1 datasets. At either threshold, Deblur achieves the best performance on the Stag1 dataset, and although AmpliCI finds one more true haplotype with threshold at 10, it is not rescued by positive filtering. It is important to note that despite Deblur’s performance on Stag1, it is far from superior on the other mock datasets and the worst performer in simulation. Additional interpretation of the comparison study on mock datasets is given in Supplementary Material S8.

**Table 1:**
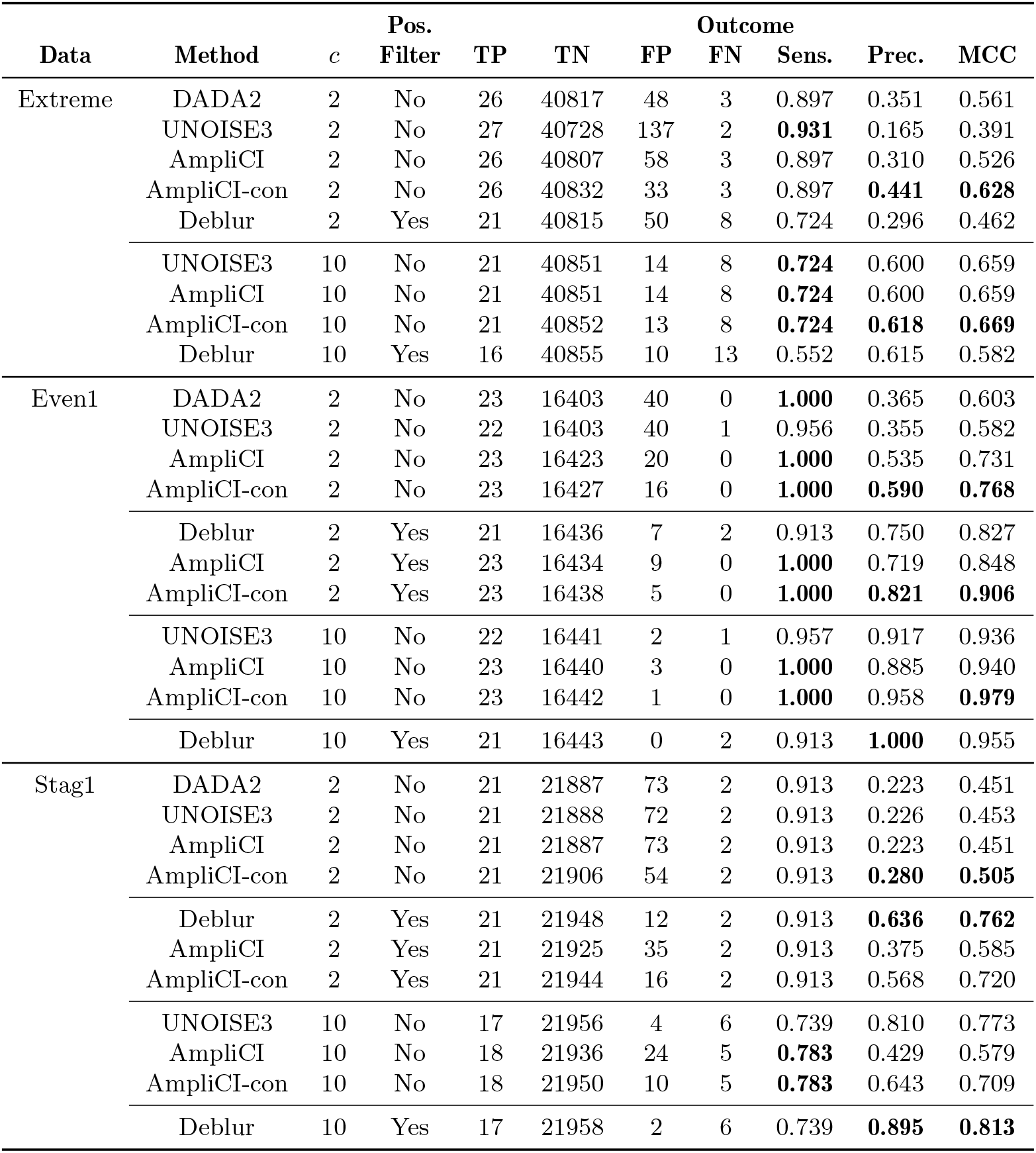
Results on Three Mock Datasets. Results on three mock datasets, Extreme, Mock5 Even1, Mock5 Stag1, treating reference sequences of the mock communities as the gold standard. Abundance threshold *c* is set as 2 or 10. AmpliCI uses default contamination diagnostics (see Methods); AmpliCI-con uses contamination diagnostic threshold 10^−40^. Since Deblur conducts a positive filtering, we also run AmpliCI with a *post hoc* positive filtering to remove likely contaminants (Pos. Filter). This positive filtering is only applied to Even1 and Stag1 datasets when *c* = 2. TP: true positives, TN: true negatives, FP: false positives, FN: false negatives, Sens.: sensitivity, Prec.: precision, MCC: Matthew’s correlation coefficient.

### 3.3. Real Dataset

The real dataset we analyze consists of 157 samples collected from 42 British women in a longitudinal study of vaginal microbiome during and after pregnancy [34]. DADA2 estimates the error profile from the first several samples until the cumulative number of nucleotides > 10^8^ and uses the same error profile to infer haplotypes in each sample independently. Then chimaera detection is performed by its default algorithm. Deblur also infers haplotypes from each sample independently with the per-sample abundance threshold at 2 (option `--min-size`) and the cross-sample abundance threshold at 10 (option `--min-reads`). UNOISE3 pools all samples together and infers haplotypes with abundance threshold at 8 (option `-minsize`). AmpliCI estimates error profiles and infers haplotypes independently for each sample. Then chimaeras are detected per sample by using UCHIME3 *de novo* [27] with default settings. We run AmpliCI with the default abundance threshold at 2 (option `-lb`), but *post hoc* filter with abundance threshold at 8 or 10 across all samples to better compare with UNOISE3 and Deblur.

Since this is a real dataset without a known reference, we evaluate the results by aligning estimated haplotypes against the Silva v132 rRNA gene database [40]. Fig. 3 shows that the total number of haplotypes with a 100% match in the database are similar among the different algorithms, although no two methods agree perfectly. Haplotypes without 100% matches in the reference database, which could be true biological variants or false positives generated during PCR and sequencing, are not inferred with as much agreement among the algorithms. Overall, AmpliCI is closest in predictions to Deblur, UNOISE3, then DADA2, although it should be noted that because more potential haplotypes are screened under the default settings for DADA2, UNOISE3, then Deblur, there are also more opportunities for disagreement in precisely the methods we find to most disagree.

**Figure 3:**
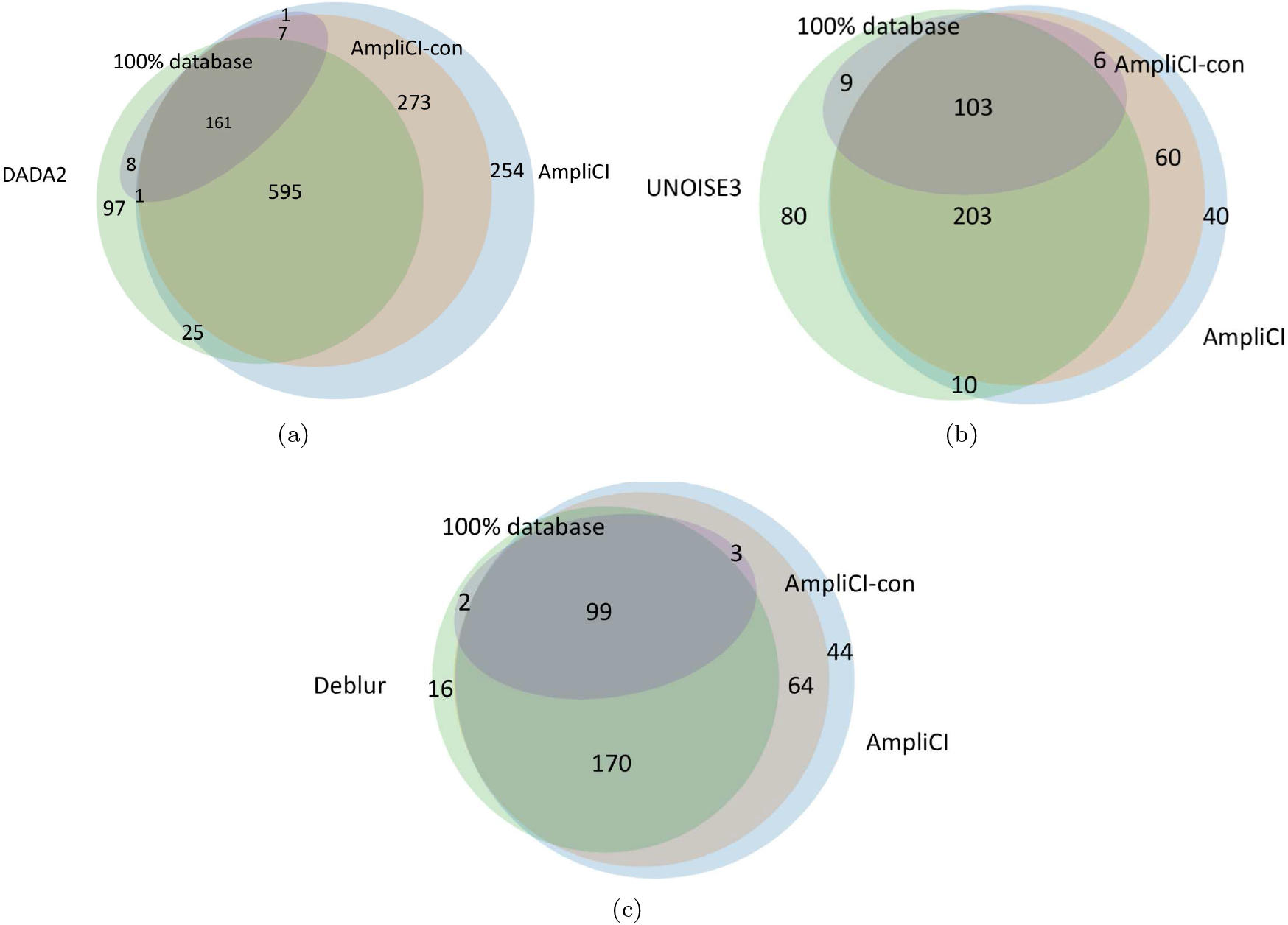
Venn diagrams of haplotypes discovered in the vaginal microbiome dataset [34] by AmpliCI, AmpliCI-con, DADA2, UNOISE3, and Deblur. Haplotypes with a 100% match to the Silva v132 rRNA gene database [40] are shaded. AmpliCI and AmpliCI-con compared to (a) DADA2 with abundance threshold *c* = 2, (b) UNOISE3 with *c* = 2 and cross-sample abundance threshold *c*\* = 8, (c) Deblur with *c* = 2 and cross-sample abundance threshold *c*\* = 10.

### 3.4. Run Time and Memory Analysis

Table 2 provides the time and memory usage of the four algorithms on the three mock and one real dataset. The GNU `time` command was used for user time and maximum memory usage on a server with an Intel(R) Xeon(R) CPU E3-1241 v3 @ 3.50GHz. For algorithms within a pipeline, we report only the timing and memory usage for the major steps of denoising. For UNOISE3 and Deblur, statistics were computed with chimaera detection via UCHIME *de novo* embedded in the denoising step, but chimaera detection is a relatively insignificant contributor to resource usage.

For datasets containing a single sample (the three mock datasets) the time and memory usage of AmpliCI increases in the number of reads and haplotypes. AmpliCI resource usage is far higher on Even1 and Stag1 than Extreme because though there are fewer reads, there are many small clusters generated by chimaeras. Though AmpliCI is not the most efficient algorithm, it uses less resources than Deblur and DADA2 on Extreme, and roughly the same resources on Even1 and Stag1. On the multi-sample vaginal microbiome, Deblur has the highest run time and DADA2 has the highest memory usage. UNOISE3 triumphs in computational and memory efficiency, beating all other methods.

**Table 2:**
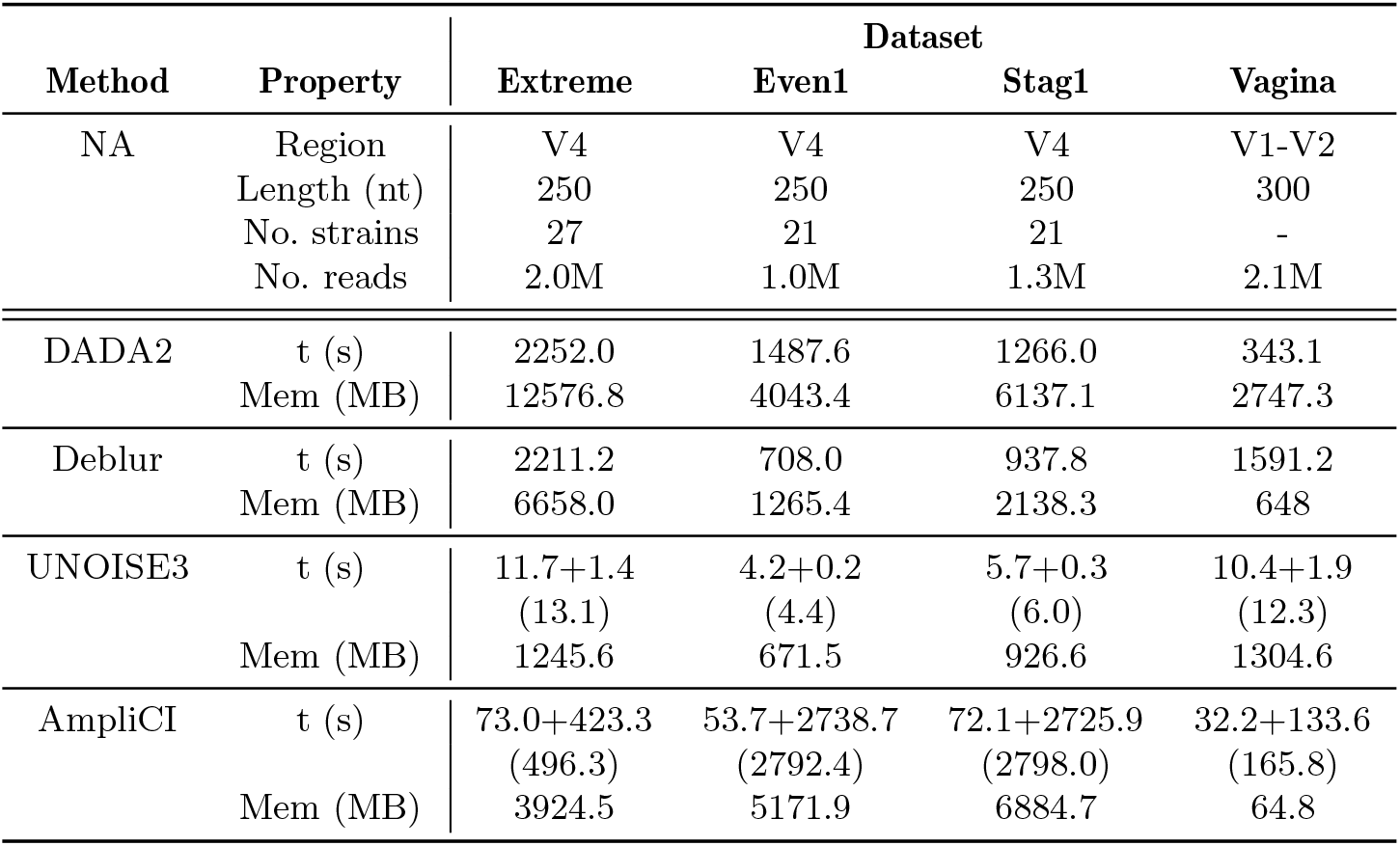
Information on Tested Datasets‥ NA: Not Applicable, Length (nt): read length in nucleotides, No. strains: number of known strains in mock community, No. reads: total number of reads in dataset, t: running time; mem: maximum resident set size, memory usage recorded by `time --verbose`. All datasets (Extreme [16], Even1 [32], Stag1 [32], Vagina [34]) contain 16S rRNA gene amplicon sequences generated from the Miseq Platform. For DADA2 and Deblur, time recorded for the whole workflow. For UNOISE3, time recorded for the two steps: data compression (-fastx uniques) and denoising (-unoise3). For AmpliCI, time recorded for the two steps: error estimation and haplotype inference. For times reported for two steps, total time reported on next line in parenthesis.

| Method | Property | Dataset |  |  |  |
| --- | --- | --- | --- | --- | --- |
|  |  | Extreme | Even1 | Stag1 | Vagina |
| NA | Region | V4 | V4 | V4 | V1-V2 |
|  | Length (nt) | 250 | 250 | 250 | 300 |
|  | No. strains | 27 | 21 | 21 | - |
|  | No. reads | 2.0M | 1.0M | 1.3M | 2.1M |
| DADA2 | t (s) | 2252.0 | 1487.6 | 1266.0 | 343.1 |
|  | Mem (MB) | 12576.8 | 4043.4 | 6137.1 | 2747.3 |
| Deblur | t (s) | 2211.2 | 708.0 | 937.8 | 1591.2 |
|  | Mem (MB) | 6658.0 | 1265.4 | 2138.3 | 648 |
| UNOISE3 | t (s) | 11.7+1.4<br>(13.1) | 4.2+0.2<br>(4.4) | 5.7+0.3<br>(6.0) | 10.4+1.9<br>(12.3) |
|  | Mem (MB) | 1245.6 | 671.5 | 926.6 | 1304.6 |
| AmpliCI | t (s) | 73.0+423.3<br>(496.3) | 53.7+2738.7<br>(2792.4) | 72.1+2725.9<br>(2798.0) | 32.2+133.6<br>(165.8) |
|  | Mem (MB) | 3924.5 | 5171.9 | 6884.7 | 64.8 |

## 4. Discussion

We propose AmpliCI, a likelihood-based denoiser of Illumina amplicon sequence data under the mixture model framework. We have shown that AmpliCI is superior in accuracy to three other popular denoising methods, and retains acceptable computation time and memory usage. Here, we discuss the advantages of AmpliCI as well as the persistent challenges (minor limitations discussed in Supplementary Material S10) that remain for all amplicon sequence denoising methods.

### 4.1. Likelihood-Based Inference with Quality Score

It is logical to formulate denoising as a clustering problem [16, 21], where all members of a cluster are reads of the same true sequence. If sequencing errors are a homogeneous disturbance of the true sequences, then the high sequencing depth supports using a homogeneous finite mixture model [41]. [4] recognized this possibility when proposing a finite mixture model for the fluorescent signals emitted by amplicon sequences processed on the 454 sequencer, but AmpliCI appears to be the first finite mixture model proposed for denoising Illumina amplicon data. AmpliCI shares several similarities with DADA2 [16], which also models errors as homogeneous disturbances on unknown true sequences, but DADA2 is not formulated as a mixture model, and the two methods take different strategies to overcome the computational challenges.

Model-based clustering is plagued by the dual computational challenges of choosing the number of clusters and global optimization, both especially challenging when sample sizes number in the millions and clusters in the hundreds to thousands. AmpliCI adopts an approximate maximization strategy for the mixture model likelihood, selecting likely true haplotypes in a greedy fashion. DADA2 does not formulate a likelihood, instead using its error model to devise a divisive clustering algorithm based on diagnostic tests. AmpliCI and DADA2 use nearly identical error models, but DADA2 compresses data at the unique sequence level, discarding the quality score information of individual reads and treating unique sequences as conditionally independent observations within clusters. In AmpliCI and as appropriate for the mixture model formulation, the *reads* are treated as the conditionally independent observations within clusters.

Using an estimated error model that considers quality scores should increase the resolution of both AmpliCI and DADA2 over other methods, such as UNOISE3 and Deblur, that use fixed thresholds and ignore quality scores. Further, AmpliCI should achieve better resolution than DADA2 by not compressing the reads into unique sequences and by capitalizing on the likelihood principle. The superiority of AmpliCI is confirmed in our simulation study, where it clearly and reproducibly detects more correct true sequences than any other method, most notably when there is little separation between the true sequences.

### 4.2. Error Models and their Limitations

Quality scores cannot be simply interpreted as Phred scores [29]. In real data, sample quality, library preparation methods, PCR conditions and sequencing platforms can generate datasets that disrupt the information communicated by Phred quality scores in sample-specific ways [24, 42, 43]. The quality information may be further invalidated by pre-denoising filters, which typically include clipping low-quality tails of reads and discarding short reads. Such filtering is recommended [44] because read quality and length can have large impact on the downstream analysis, especially diversity estimation [45]. Simply converting quality scores to Phred error probabilities can indeed be disastrous: AmpliCI is overrun with haplotype predictions (858 compared to 29 when using estimated errors with abundance threshold *c* = 15 on the Extreme dataset).

The error models inside Deblur and UNOISE3 are fixed across samples, ignore the differences in substitution miscalls (transitions A ↔ G and T ↔ C are usually more common than transversions), and ignore quality scores all together [18, 24, 42]. While DADA2 and AmpliCI estimate and use quality score-based error models, they do not account for all the patterns in sequencing errors. Systematic sequencing errors have been identified in Illumina data [46, 24, 42], including high error rates that associate with certain 3-mer motifs and inverted repeats [24, 46]. Even worse, real data are replete with additional false signals (contaminants, propagated PCR errors) that mimic true haplotypes.

Methods can handle this complexity by (1) modeling it, (2) assuming error rates high enough to bury it in the noise, (3) discarding low abundance variants, or (4) making conservative decisions. We are aware of no method that has successfully done (1), and all remaining strategies sacrifice resolving power. UNOISE3 and Deblur adopt strategies (2) and (3) by setting conservative error rates and recommending high abundance thresholds. Less obviously, but to lesser degree, both AmpliCI and DADA2 also adopt (2) and (3). Neither can consider singletons as possible haplotypes, and because they use an approximate strategy to estimate error rates *before* all haplotypes have been determined, the error rates are overestimated. A better strategy is to invent better error models (1), while applying conservative decisions (4) to overcome remaining deficiencies. A careful comparison of method performance on all three mock datasets, shows that AmpliCI has achieved a better trade-off in decision errors by more accurately modeling the per-read sequencing process. Higher resolution denoisers will be achieved through better understanding and models for the as yet untackled noise in amplicon sequencing.

## Acknowledgements

We thank Heliang Shi for previous work on *Ampliclust*, an EM algorithm for maximization of Eq. (1). An earlier version of this paper won the American Statistical Association Section on Statistics in Genomics and Genetics’ 2020 Student Paper Competition.

## References

[1] R. Knight, A. Vrbanac, B. C. Taylor, A. Aksenov, C. Callewaert, J. Debelius, A. Gonzalez, T. Kosci-olek, L.-I. McCall, D. McDonald, A. V. Melnik, J. T. Morton, J. Navas, R. A. Quinn, J. G. Sanders, A. D. Swafford, L. R. Thompson, A. Tripathi, Z. Z. Xu, J. R. Zaneveld, Q. Zhu, J. G. Caporaso, P. C. Dorrestein, Best practices for analysing microbiomes, Nature Reviews Microbiology 16 (2018) 1–13.

[2] B. J. Callahan, P. J. McMurdie, S. P. Holmes, Exact sequence variants should replace operational taxonomic units in marker-gene data analysis, The ISME Journal 11 (2017) 2639–2643.

[3] R. C. Edgar, Accuracy of microbial community diversity estimated by closed- and open-reference OTUs, PeerJ 5 (2017) e3889.

[4] C. Quince, A. Lanzén, T. P. Curtis, R. J. Davenport, N. Hall, I. M. Head, L. F. Read, W. T. Sloan, Accurate determination of microbial diversity from 454 pyrosequencing data, Nature Methods 6 (2009) 639–641.

[5] S. M. Huse, J. A. Huber, H. G. Morrison, M. L. Sogin, D. M. Welch, Accuracy and quality of massively parallel DNA pyrosequencing, Genome Biology 8 (2007) R143.

[6] S. M. Huse, D. M. Welch, H. G. Morrison, M. L. Sogin, Ironing out the wrinkles in the rare biosphere through improved OTU clustering, Environmental Microbiology 12 (2010) 1889–1898.

[7] E. Kopylova, J. A. Navas-Molina, C. Mercier, Z. Z. Xu, F. Mahé, Y. He, H.-W. Zhou, T. Rognes, J. G. Caporaso, R. Knight, Open-source sequence clustering methods improve the state of the art, mSystems 1 (2016) e00003–15.

[8] J. T. Nearing, G. M. Douglas, A. M. Comeau, M. G. I. Langille, J. Chen, Denoising the denoisers: an independent evaluation of microbiome sequence error-correction approaches, PeerJ 6 (2018) e5364.

[9] K. T. Konstantinidis, J. M. Tiedje, Genomic insights that advance the species definition for prokaryotes, Proceedings of the National Academy of Sciences 102 (2005) 2567–2572.

[10] E. Stackebrandt, B. M. Goebel, Taxonomic note: A place for DNA-DNA reassociation and 16S rRNA sequence analysis in the present species definition in bacteriology, International Journal of Systematic and Evolutionary Microbiology 44 (1994) 846–849.

[11] P. D. Schloss, S. L. Westcott, Assessing and improving methods used in operational taxonomic unit-based approaches for 16S rRNA gene sequence analysis, Applied and Environmental Microbiology 77 (2011) 3219–3226.

[12] R. C. Edgar, Updating the 97% identity threshold for 16S ribosomal RNA OTUs, Bioinformatics 34 (2018) 2371–2375.

[13] E. Stackebrandt, J. Ebers, Taxonomic parameters revisited: tarnished gold standards, Microbiology Today 33 (2006) 152–155.

[14] J. S. Johnson, D. J. Spakowicz, B.-Y. Hong, L. M. Petersen, P. Demkowicz, L. Chen, S. R. Leopold, B. M. Hanson, H. O. Agresta, M. Gerstein, E. Sodergren, G. M. Weinstock, Evaluation of 16S rRNA gene sequencing for species and strain-level microbiome analysis, Nature Communications 10 (2019) 5029.

[15] M. Rossi-Tamisier, S. Benamar, D. Raoult, P.-E. Fournier, Cautionary tale of using 16S rRNA gene sequence similarity values in identification of human-associated bacterial species, International Journal of Systematic and Evolutionary Microbiology 65 (2015) 1929–1934.

[16] B. J. Callahan, P. J. McMurdie, M. J. Rosen, A. W. Han, A. J. A. Johnson, S. P. Holmes, DADA2: High-resolution sample inference from Illumina amplicon data., Nature Methods 13 (2016) 581–583.

[17] A. M. Eren, L. Maignien, W. J. Sul, L. G. Murphy, S. L. Grim, H. G. Morrison, M. L. Sogin, Oligotyping: Differentiating between closely related microbial taxa using 16S rRNA gene data, Methods in Ecology and Evolution 4 (2013) 1111–1119.

[18] M. Tikhonov, R. W. Leach, N. S. Wingreen, Interpreting 16S metagenomic data without clustering to achieve sub-OTU resolution, The ISME Journal 9 (2014) 68.

[19] A. M. Eren, H. G. Morrison, P. J. Lescault, J. Reveillaud, J. H. Vineis, M. L. Sogin, Minimum entropy decomposition: Unsupervised oligotyping for sensitive partitioning of high-throughput marker gene sequences, The ISME Journal 9 (2015) 968–979.

[20] M. Mysara, N. Leys, J. Raes, P. Monsieurs, IPED: a highly efficient denoising tool for Illumina MiSeq paired-end 16S rRNA gene amplicon sequencing data, BMC Bioinformatics 17 (2016) 192.

[21] R. C. Edgar, UNOISE2: improved error-correction for Illumina 16S and ITS amplicon sequencing, bioRxiv 081257.

[22] A. Amir, D. McDonald, J. A. Navas-Molina, E. Kopylova, J. T. Morton, Z. Zech Xu, E. P. Kightley, L. R. Thompson, E. R. Hyde, A. Gonzalez, R. Knight, Deblur rapidly resolves single-nucleotide community sequence patterns, mSystems 2 (2017) e00191–16.

[23] N. J. Hathaway, C. M. Parobek, J. J. Juliano, J. A. Bailey, SeekDeep: single-base resolution de novo clustering for amplicon deep sequencing, Nucleic Acids Research 46 (2017) e21.

[24] M. Schirmer, U. Z. Ijaz, R. D’Amore, N. Hall, W. T. Sloan, C. Quince, Insight into biases and sequencing errors for amplicon sequencing with the Illumina MiSeq platform, Nucleic Acids Research 43 (2015) e37.

[25] V. Melnykov, R. Maitra, Finite mixture models and model-based clustering, Statistics Surveys 4 (2010) 80–116.

[26] X. Yang, S. Aluru, K. S. Dorman, Repeat-aware modeling and correction of short read errors., BMC Bioinformatics 12 (2011) S52.

[27] R. Edgar, UCHIME2: improved chimera prediction for amplicon sequencing, bioRxiv 74252.

[28] A. McKenna, M. Hanna, E. Banks, A. Sivachenko, K. Cibulskis, A. Kernytsky, K. Garimella, D. Altshuler, S. Gabriel, M. Daly, M. A. DePristo, The Genome Analysis Toolkit: A MapReduce framework for analyzing next-generation DNA sequencing data, Genome Research 20 (2010) 1297–1303.

[29] B. Ewing, P. Green, Base-calling of automated sequencer traces using Phred. II. error probabilities, Genome Research 8 (1998) 186–194.

[30] W. Huang, L. Li, J. R. Myers, G. T. Marth, ART: a next-generation sequencing read simulator, Bioinformatics 28 (2012) 593–594.

[31] T. H. Jukes, C. R. Cantor, Evolution of protein molecules, in: H. N. Munro, J. B. Allison (Eds.), Mammalian Protein Metabolism, Vol. 3, Academic Press, New York, 1969, pp. 21–132.

[32] N. A. Bokulich, J. R. Rideout, E. Kopylova, E. Bolyen, J. Patnode, Z. Ellett, D. McDonald, B. Wolfe, C. F. Maurice, R. J., A standardized, extensible framework for optimizing classification improves marker-gene taxonomic assignments, PeerJ PrePrints 3 (2015) e934v2.

[33] N. A. Bokulich, J. R. Rideout, W. G. Mercurio, A. Shiffer, B. Wolfe, C. F. Maurice, R. J. Dutton, P. J. Turnbaugh, R. Knight, J. G. Caporaso, mockrobiota: a public resource for microbiome bioinformatics benchmarking, mSystems 1 (2016) e00062–16.

[34] D. A. MacIntyre, M. Chandiramani, Y. S. Lee, L. Kindinger, A. Smith, N. Angelopoulos, B. Lehne, S. Arulkumaran, R. Brown, T. G. Teoh, E. Holmes, J. K. Nicoholson, J. R. Marchesi, P. R. Bennett, The vaginal microbiome during pregnancy and the postpartum period in a European population, Scientific Reports 5 (2015) 8988.

[35] J. G. Caporaso, J. Kuczynski, J. Stombaugh, K. Bittinger, F. D. Bushman, E. K. Costello, N. Fierer, A. G. Peña, J. K. Goodrich, J. I. Gordon, G. A. Huttley, S. T. Kelley, D. Knights, J. E. Koenig, R. E. Ley, C. A. Lozupone, D. McDonald, B. D. Muegge, M. Pirrung, J. Reeder, J. R. Sevinsky, P. J. Turnbaugh, W. A. Walters, J. Widmann, T. Yatsunenko, J. Zaneveld, R. Knight, QIIME allows analysis of high-throughput community sequencing data, Nature Methods 7 (2010) 335–336.

[36] T. Z. DeSantis, P. Hugenholtz, N. Larsen, M. Rojas, E. L. Brodie, K. Keller, T. Huber, D. Dalevi, P. Hu, G. L. Andersen, Greengenes, a chimera-checked 16S rRNA gene database and workbench compatible with ARB, Applied and Environmental Microbiology 72 (2006) 5069–5072.

[37] E. Kopylova, L. Noé, H. Touzet, SortmeRNA: fast and accurate filtering of ribosomal RNAs in metatranscriptomic data, Bioinformatics 28 (2012) 3211–3217.

[38] R. C. Edgar, Search and clustering orders of magnitude faster than BLAST, Bioinformatics 26 (2010) 2460–2461.

[39] L. Hubert, P. Arabie, Comparing partitions, Journal of Classification 2 (1985) 193–218.

[40] P. Yilmaz, L. W. Parfrey, P. Yarza, J. Gerken, E. Pruesse, C. Quast, T. Schweer, J. Peplies, W. Ludwig, F. O. Glöckner, The SILVA and “all-species living tree project (LTP)” taxonomic frameworks, Nucleic Acids Research 42 (2014) D643–D648.

[41] G. J. McLachlan, D. Peel, Finite Mixture Models, Wiley series in probability and statistics, Wiley, New York, 2000.

[42] M. Schirmer, U. Z. Ijaz, N. Hall, C. Quince, Illumina error profiles: Resolving fine-scale variation in metagenomic sequencing data, BMC Bioinformatics 17 (2016) 1–15.

[43] J. M. Bender, F. Li, H. Adisetiyo, D. Lee, S. Zabih, L. Hung, T. A. Wilkinson, P. S. Pannaraj, R. C. She, J. D. Bard, N. H. Tobin, G. M. Aldrovandi, Quantification of variation and the impact of biomass in targeted 16S rRNA gene sequencing studies, Microbiome 6 (2018) 155.

[44] J. G. Caporaso, C. L. Lauber, W. A. Walters, D. Berg-Lyons, C. A. Lozupone, P. J. Turnbaugh, N. Fierer, R. Knight, Global patterns of 16S rRNA diversity at a depth of millions of sequences per sample, Proceedings of the National Academy of Sciences 108 (2011) 4516–4522.

[45] N. A. Bokulich, S. Subramanian, J. J. Faith, D. Gevers, J. I. Gordon, R. Knight, D. A. Mills, J. G. Caporaso, Quality-filtering vastly improves diversity estimates from Illumina amplicon sequencing, Nature Methods 10 (2012) 57–59.

[46] K. Nakamura, T. Oshima, T. Morimoto, S. Ikeda, H. Yoshikawa, Y. Shiwa, S. Ishikawa, M. C. Linak, A. Hirai, H. Takahashi, M. Altaf-Ul-Amin, N. Ogasawara, S. Kanaya, Sequence-specific error profile of Illumina sequencers, Nucleic Acids Research 39 (2011) e90–e90.

[47] S. F. Altschul, W. Gish, W. Miller, E. W. Myers, D. J. Lipman, Basic local alignment search tool, Journal of Molecular Biology 215 (1990) 403–410.

[48] P. Yilmaz, L. W. Parfrey, P. Yarza, J. Gerken, E. Pruesse, C. Quast, T. Schweer, J. Peplies, W. Ludwig, F. O. Glöckner, The SILVA and “All-species Living Tree Project (LTP)” taxonomic frameworks, Nucleic Acids Research 42 (2013) D643–D648.

[49] M. M. Weinstein, A. Prem, M. Jin, S. Tang, J. M. Bhasin, FIGARO: An efficient and objective tool for optimizing microbiome rRNA gene trimming parameters, bioRxiv 610394.

[50] R. Edgar, Taxonomy annotation and guide tree errors in 16S rRNA databases, PeerJ 6 (2018) e5030.

[51] J. Ritari, J. Salojärvi, L. Lahti, W. M. de Vos, Improved taxonomic assignment of human intestinal 16S rRNA sequences by a dedicated reference database, BMC Genomics 16 (2015) 1056.

[52] G. Kallós, A generalization of Pascal’s triangle using powers of base numbers, Annales Mathématiques Blaise Pascal 13 (2006) 1–15.

[53] Y. Hong, On computing the distribution function for the Poisson binomial distribution, Computational Statistics and Data Analysis 59 (2013) 41–51.

[54] S. J. Salter, M. J. Cox, E. M. Turek, S. T. Calus, W. O. Cookson, M. F. Moffatt, P. Turner, J. Parkhill, N. J. Loman, A. W. Walker, Reagent and laboratory contamination can critically impact sequence-based microbiome analyses, BMC Biology 12 (2014) 87–87.

[55] G. Renaud, U. Stenzel, J. Kelso, leeHom: adaptor trimming and merging for Illumina sequencing reads., Nucleic Acids Research 42 (2014) e141.

