## Supplemental Materials for "AmpliCI: A High-resolution Model-Based Approach for Denoising Illumina Amplicon Data"

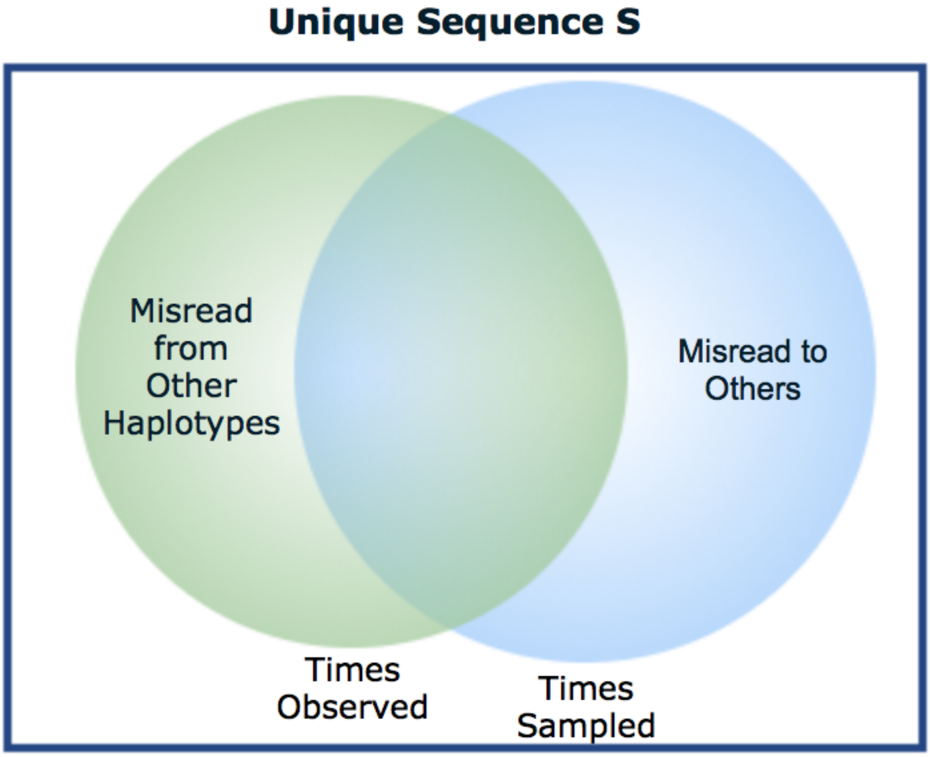


Figure S1: The core idea behind AmpliCI. The goal is to estimate the true abundance of each observed unique sequence $\boldsymbol{s}$ (area of blue circle). Unfortunately, the number of times $\boldsymbol{s}$ is observed (area of green circle) includes misreads from other unique sequences and excludes misreads of $\boldsymbol{s}$, both due to errors during PCR and sequencing. Thus, information about the green area does not readily translate into information about the blue area. However, if the misread rate is constant for all haplotypes, we can estimate the total number of reads in the overlap region, which is directly proportional to the blue area and true abundance. We call the resulting quantity the *estimated scaled true abundance* and use it to populate Amplicon Sequence Variant (ASV) abundance tables.

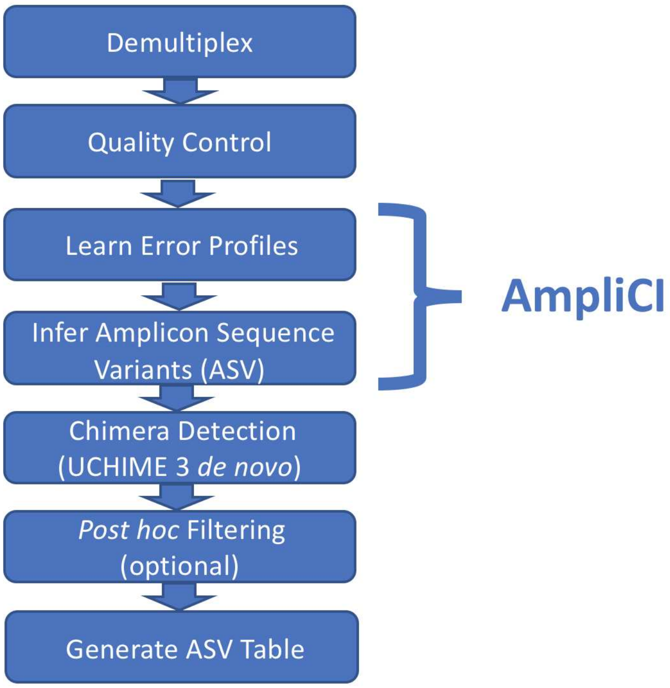


Figure S2: Analysis pipeline used for denoising amplicon sequence data. Starting with one or more samples of Illumina amplicon sequence data in FASTQ format, forward reads are demultiplexed and filtered for quality and length. AmpliCI is run once to estimate error rates for each sample, then again with the estimated error rates to infer Amplicon Sequence Variants (ASVs) and abundance for each sample. UCHIME3 *de novo*[1] is applied for chimaera detection. More conservative haplotype predictions are optionally achieved by applying additional *post hoc* filtering, for example to apply more stringent thresholds on diagnostics or estimated abundances. The output is an ASV table recording ASVs and their abundances in each sample.

| $c_d$ | Mean Similarity | AmpliCI K(T.seq) | AmpliCI ARI | DADA2 K(T.seq) | DADA2 ARI | UNOISE3 K(T.seq) | UNOISE3 ARI | Deblur K(T.seq) | Deblur ARI |
|---|---|---|---|---|---|---|---|---|---|
| | 0.986 | **10(10)** | **0.998** | 9(9) | 0.997 | 11(10) | 0.997 | 4(4) | 0.869 |
| 0.01 | 0.983 | **11(11)** | **0.986** | 9(9) | 0.968 | 7(7) | 0.726 | 4(4) | 0.623 |
| | 0.989 | **10(10)** | **0.979** | 8(8) | 0.959 | 7(7) | 0.904 | 4(4) | 0.670 |
| | 0.975 | **12(12)** | **1.000** | 9(9) | 0.981 | 13(12) | 1.000 | 7(7) | 0.899 |
| 0.02 | 0.976 | **11(11)** | **0.995** | 9(9) | 0.987 | 10(10) | 0.988 | 7(7) | 0.925 |
| | 0.974 | **12(12)** | **0.999** | 10(10) | 0.990 | 12(12) | 0.997 | 7(7) | 0.914 |
| | 0.966 | **12(12)** | **1.000** | 10(10) | 0.998 | 13(12) | 1.000 | 8(8) | 0.996 |
| 0.03 | 0.966 | **12(12)** | **1.000** | 10(10) | 0.988 | 11(11) | 0.993 | 8(8) | 0.984 |
| | 0.969 | **12(12)** | **1.000** | 11(11) | 0.995 | 12(12) | 0.999 | 7(7) | 0.923 |
| | 0.962 | **12(12)** | **1.000** | 10(10) | 0.999 | 13(12) | 1.000 | 9(9) | 0.999 |
| 0.04 | 0.958 | 12(12) | 1.000 | 11(11) | 0.996 | 12(12) | 1.000 | 9(9) | 0.990 |
| | 0.963 | 12(12) | 1.000 | 11(11) | 0.999 | 12(12) | 1.000 | 9(9) | 0.997 |
| | 0.954 | **12(12)** | **1.000** | 11(11) | 0.996 | 13(12) | 1.000 | 10(10) | 0.995 |
| 0.05 | 0.952 | 12(12) | 1.000 | 12(12) | 1.000 | 12(12) | 1.000 | 10(10) | 0.992 |
| | 0.950 | 12(12) | 1.000 | 12(12) | 1.000 | 12(12) | 1.000 | 11(11) | 1.000 |
| | 0.799 | 11(11) | 0.999 | 11(11) | 0.999 | 11(11) | 1.000 | 11(11) | 1.000 |
| - | 0.799 | 12(12) | 1.000 | 12(12) | 1.000 | 14(12) | 0.999 | 12(12) | 0.999 |
| | 0.799 | 12(12) | 1.000 | 12(12) | 1.000 | 12(12) | 1.000 | 12(12) | 1.000 |

Table S1: Simulation results. All three replicate datasets per simulation condition contain 3,000 reads generated from 12 true haplotypes. $c_d$ is the evolutionary distance, in the Jukes-Cantor model, controlling the similarities between synthetic haplotypes. Datasets with $c_d = -$ are simulated using 12 real 16S rRNA gene V4 sequences from the Extreme dataset as true haplotypes. The lowest allowed observed abundance of haplotypes is set as 2 for all methods. $K$: total number of haplotypes identified, T.seq: number of true haplotypes identified, ARI: Adjusted Rand Index. Bolded numbers in each row indicate the best performer unless there is a tie.

| Datasets | Methods | c | Pos. Filter | Total | Expected 100% | Expected 97% | Database 100% | Database 97% | Unmatched |
|---|---|---|---|---|---|---|---|---|---|
| Extreme | DADA2 | 2 | No | 74 | 26 | 8 | 12 | 2 | 26 |
| | UNOISE3 | 2 | No | 164 | **27** | 21 | 13 | 4 | 99 |
| | AmpliCI | 2 | No | 84 | 26 | 3 | 12 | 3 | 40 |
| | AmpliCI-con | 2 | No | 59 | 26 | **1** | 9 | 2 | 21 |
| | Deblur | 2 | Yes | 71 | 21 | 3 | 12 | 4 | 31 |
| | UNOISE3 | 10 | No | 35 | **21** | 1 | 2 | 1 | 10 |
| | AmpliCI | 10 | No | 35 | **21** | 1 | 2 | 1 | 10 |
| | AmpliCI-con | 10 | No | 34 | **21** | 1 | 2 | 1 | 9 |
| | Deblur | 10 | Yes | 26 | 16 | **0** | 2 | 1 | 7 |
| Even1 | DADA2 | 2 | No | 63 | **23** | 25 | 1 | 1 | 13 |
| | UNOISE3 | 2 | No | 62 | 22 | 14 | 1 | 4 | 21 |
| | AmpliCI | 2 | No | 43 | **23** | 4 | 1 | 2 | 13 |
| | AmpliCI-con | 2 | No | 39 | **23** | **1** | 1 | 1 | 13 |
| | Deblur | 2 | Yes | 28 | 21 | **1** | 1 | 1 | 4 |
| | AmpliCI | 2 | Yes | 32 | **23** | 4 | 1 | 1 | 2 |
| | AmpliCI-con | 2 | Yes | 28 | **23** | **1** | 1 | 1 | 2 |
| | UNOISE3 | 10 | No | 24 | 22 | 2 | 0 | 0 | 0 |
| | AmpliCI | 10 | No | 26 | **23** | 3 | 0 | 0 | 0 |
| | AmpliCI-con | 10 | No | 24 | **23** | 1 | 0 | 0 | 0 |
| | Deblur | 10 | Yes | 21 | 21 | **0** | 0 | 0 | 0 |
| Stag1 | DADA2 | 2 | No | 94 | 21 | 26 | 3 | 2 | 42 |
| | UNOISE3 | 2 | No | 93 | 21 | 17 | 4 | 5 | 46 |
| | AmpliCI | 2 | No | 94 | 21 | 27 | 3 | 2 | 41 |
| | AmpliCI-con | 2 | No | 75 | 21 | **9** | 3 | 2 | 40 |
| | Deblur | 2 | Yes | 33 | 21 | **0** | 3 | 1 | 8 |
| | AmpliCI | 2 | Yes | 56 | 21 | 19 | 3 | 2 | 11 |
| | AmpliCI-con | 2 | Yes | 37 | 21 | 5 | 3 | 2 | 6 |
| | UNOISE3 | 10 | No | 21 | 17 | 1 | 1 | 2 | 0 |
| | AmpliCI | 10 | No | 42 | **18** | 21 | 1 | 2 | 0 |
| | AmpliCI-con | 10 | No | 28 | **18** | 7 | 1 | 2 | 0 |
| | Deblur | 10 | Yes | 19 | 17 | **0** | 1 | 1 | 0 |

Table S2: Results on three mock datasets, Extreme, Mock5 Even1, and Mock5 Stag1, after aligning results to mock reference sequences and the Silva v132 rRNA gene database [2] using BLASTN v2.8.1+[3]. Expected 100%: Number of haplotypes with 100% match to the mock reference sequence. Expected 97%: Number of haplotypes without 100% match, but >97% similarity to the mock reference sequences. The remaining sequences with similarities <97% to mock references are aligned to the Silva v132 database [2]. Database 100%: Number of remaining haplotypes with 100% match to the database. Database 97%: Number of remaining haplotypes without 100% match, but >97% similarity to the database. The best performance in columns Expected 100%, and Expected 97% are bolded. Other columns are described in the Table 1 legend in the main text.

| Method | Merged Reads | Pooled | Error Estimation | Pos. Filter | Open Source | Implemen-tation | Version Tested | Refer-ence |
|---|---|---|---|---|---|---|---|---|
| DADA2 | No | No | Yes | No | Yes | R | v1.8.0 | [4] |
| UNOISE3 | Yes | Yes | No | No | No | C++ | v11.0 | [5] |
| Deblur | No | No | No | Yes | Yes | Python | v1.1.0 | [6] |
| AmpliCI | No | No | Yes | No | Yes | C | v1.0 | - |

Table S3: Comparison of AmpliCI, DADA2, UNOISE3 and Deblur. Merged Reads: whether the method recommends merging reads before denoising. Pooled: whether the method recommends pooling samples before denoising. Pos. Filter: whether reads are positively filtered against a 16S rRNA database. For UNOISE3, the 32-bit free academic version of USEARCH, version 11, was used in this study.

#### S1 Derivation of the Abundance Estimation Function

Haplotypes are selected based on the estimated scaled abundance

$$\tilde{a}^{(k)}_{\boldsymbol{s}_m} := a_{\boldsymbol{s}_m o} - \mathbb{E}[N_{\mathcal{H}_k \boldsymbol{s}_m} \mid \mathcal{H}_k; \mathcal{Q}_m].$$

To compute $\mathbb{E}[N_{\mathcal{H}_k \boldsymbol{s}_m} \mid \mathcal{H}_k; \mathcal{Q}_m]$, we let $\boldsymbol{Z}_{mi}$ (reindexed) represent the haplotype source of the $i$th read $\boldsymbol{R}_{mi}$ matching the $m$th unique sequence. If the only possible sources of reads with sequence $\boldsymbol{s}_m$ are the current set of haplotypes $\mathcal{H}_k$ or $\boldsymbol{s}_m$ itself, then the expected number of misreads is

$$\begin{aligned}
\mathbb{E}[N_{\mathcal{H}_k \boldsymbol{s}_m} \mid \mathcal{H}_k; \mathcal{Q}_m] &= \sum_{i=1}^{a_{\boldsymbol{s}_m o}} \Pr(\boldsymbol{Z}_{mi} \in \mathcal{H}_k \mid \boldsymbol{R}_{mi} = \boldsymbol{s}_m, \boldsymbol{Z}_{mi} \in \{\mathcal{H}_k, \boldsymbol{s}_m\}; \boldsymbol{q}_{mi}) \\
&= \sum_{i=1}^{a_{\boldsymbol{s}_m o}} \sum_{w=1}^{|\mathcal{H}_k|} \Pr(\boldsymbol{Z}_{mi} = \boldsymbol{h}_w \mid \boldsymbol{R}_{mi} = \boldsymbol{s}_m, \boldsymbol{Z}_{mi} \in \{\mathcal{H}_k, \boldsymbol{s}_m\}; \boldsymbol{q}_{mi}) \\
&= \sum_{i=1}^{a_{\boldsymbol{s}_m o}} \sum_{w=1}^{|\mathcal{H}_k|} \frac{\Pr(\boldsymbol{R}_{mi} = \boldsymbol{s}_m \mid \boldsymbol{Z}_{mi} = \boldsymbol{h}_w; \boldsymbol{q}_{mi}) \Pr(\boldsymbol{Z}_{mi} = \boldsymbol{h}_w \mid \boldsymbol{Z}_{mi} \in \{\mathcal{H}_k, \boldsymbol{s}_m\})}{\Pr(\boldsymbol{R}_{mi} = \boldsymbol{s}_m \mid \boldsymbol{Z}_{mi} \in \{\mathcal{H}_k, \boldsymbol{s}_m\}; \boldsymbol{q}_{mi})}
\end{aligned}$$

where

$$\begin{aligned}
&\Pr(\boldsymbol{R}_{mi} = \boldsymbol{s}_m \mid \boldsymbol{Z}_{mi} \in \{\mathcal{H}_k, \boldsymbol{s}_m\}; \boldsymbol{q}_{mi}) \\
&\quad = \sum_{w=1}^{|\mathcal{H}_k|} \Pr(\boldsymbol{R}_{mi} = \boldsymbol{s}_m \mid \boldsymbol{Z}_{mi} = \boldsymbol{h}_w; \boldsymbol{q}_{mi}) \Pr(\boldsymbol{Z}_{mi} = \boldsymbol{h}_w \mid \boldsymbol{Z}_{mi} \in \{\mathcal{H}_k, \boldsymbol{s}_m\}) \\
&\qquad + \Pr(\boldsymbol{R}_{mi} = \boldsymbol{s}_m \mid \boldsymbol{Z}_{mi} = \boldsymbol{s}_m; \boldsymbol{q}_{mi}) \Pr(\boldsymbol{Z}_{mi} = \boldsymbol{s}_m \mid \boldsymbol{Z}_{mi} \in \{\mathcal{H}_k, \boldsymbol{s}_m\})
\end{aligned}$$

and $\Pr(\boldsymbol{R}_{mi} = \boldsymbol{s}_m \mid \boldsymbol{Z}_{mi} = \boldsymbol{s}; \boldsymbol{q}_{mi})$ (succinctly $e_{\boldsymbol{s}mi}$) is the probability that a copy of any real sequence $\boldsymbol{s}$ in the current haplotype set $\{\mathcal{H}_k, \boldsymbol{s}_m\}$ is sequenced as read sequence $\boldsymbol{s}_m$, given quality scores $\boldsymbol{q}_{mi}$.

If $b = \sum_{\boldsymbol{h} \in \mathcal{H}_k} \tilde{a}_{\boldsymbol{h}} + \tilde{a}_{\boldsymbol{s}_m}$ is the sum of scaled abundances in the presumed haplotype set $\{\mathcal{H}_k, \boldsymbol{s}_m\}$, then we may estimate

$$\Pr(\boldsymbol{Z}_{mi} = \boldsymbol{s} \mid \boldsymbol{Z}_{mi} \in \{\mathcal{H}_k, \boldsymbol{s}_m\}) = \frac{\tilde{a}_{\boldsymbol{s}}}{b},$$

for $\boldsymbol{s} \in \mathcal{H}_k$. Constant $b$ is unknown because $\tilde{a}_{\boldsymbol{s}_m}$ is unknown, but it cancels in the estimation formula, which becomes

$$\tilde{a}^{(k)}_{\boldsymbol{s}_m} = a_{\boldsymbol{s}_m o} - \sum_{i=1}^{a_{\boldsymbol{s}_m o}} \sum_{w=1}^{|\mathcal{H}_k|} \frac{e_{\boldsymbol{h}_w mi} \tilde{a}_{\boldsymbol{h}_w}}{\sum_{l=1}^{|\mathcal{H}_k|} e_{\boldsymbol{h}_l mi} \tilde{a}_{\boldsymbol{h}_l} + e_{\boldsymbol{s}_m mi} \tilde{a}^{(k)}_{\boldsymbol{s}_m}}. \tag{S1}$$

#### S2 Solution of the Updating Equation

Our goal is to solve the updating equation (S1) (Eq. (5) in the main text) to estimate $\tilde{a}_{\boldsymbol{s}_m}^{(k)}$. Since the second term in Eq. (S1) is positive and bounded above by $a_{\boldsymbol{s}_m o}$, the fixed point iterator maps the interval $[0, a_{\boldsymbol{s}_m o}]$ to itself. Thus, there exists at least one fixed point in the interval by the Brouwer fixed-point theorem. Clearly, $\tilde{a}_{\boldsymbol{s}_m}^{(k)} = 0$ is a solution of the equation. Here, we investigate possible conditions for a positive solution $\tilde{a}_{\boldsymbol{s}_m}^{(k)} > 0$. Let $x = \tilde{a}_{\boldsymbol{s}_m}^{(k)}$ and

$$c_i = \frac{e_{\boldsymbol{s}_m mi}}{\sum_{w=1}^{|\mathcal{H}_k|} e_{\boldsymbol{h}_w mi} \tilde{a}_{\boldsymbol{h}_w}} > 0, \ i = 1, 2, \cdots, a_{\boldsymbol{s}_m o}.$$

Then rewrite the equation as

$$\frac{x}{a_{\boldsymbol{s}_m o}} = 1 - \frac{1}{a_{\boldsymbol{s}_m o}} \sum_{i=1}^{a_{\boldsymbol{s}_m o}} \frac{1}{1 + c_i x} \leq 1 - \frac{1}{1 + \bar{c} x},$$

where we have applied Jensen's inequality to the convex function $\varphi(c) = \frac{1}{1+cx}$ with $a_{\boldsymbol{s}_m o} \bar{c} = \sum_{i=1}^{a_{\boldsymbol{s}_m o}} c_i$. If $x \geq 0$, then $\bar{c} x \leq \bar{c} a_{\boldsymbol{s}_m o} - 1$, thus a necessary condition for a positive solution of (S1) is

$$\bar{c} a_{\boldsymbol{s}_m o} > 1.$$

#### S3 Convergence of the Fixed Point Iterator

If $\bar{c} a_{\boldsymbol{s}_m o} \leq 1$, we know $\tilde{a}_{\boldsymbol{s}_m}^{(k)} = 0$. Otherwise, if $f(\cdot)$ is the fixed point iterator of the updating equation (S1) (Eq. (5) in the main text) with fixed point $\tilde{a}_{\boldsymbol{s}_m}^{(k)}$ and $x > \tilde{a}_{\boldsymbol{s}_m}^{(k)}$, then

$$\sum_{i=1}^{a_{\boldsymbol{s}_m o}} \sum_{w=1}^{|\mathcal{H}_k|} \frac{e_{\boldsymbol{h}_w mi} \tilde{a}_{\boldsymbol{h}_w}}{\sum_{l=1}^{|\mathcal{H}_k|} e_{\boldsymbol{h}_l mi} \tilde{a}_{\boldsymbol{h}_l} + e_{\boldsymbol{s}_m mi} x} < \sum_{i=1}^{a_{\boldsymbol{s}_m o}} \sum_{w=1}^{|\mathcal{H}_k|} \frac{e_{\boldsymbol{h}_w mi} \tilde{a}_{\boldsymbol{h}_w}}{\sum_{l=1}^{|\mathcal{H}_k|} e_{\boldsymbol{h}_l mi} \tilde{a}_{\boldsymbol{h}_l} + e_{\boldsymbol{s}_m mi} \tilde{a}_{\boldsymbol{s}_m}^{(k)}},$$

and the next iterate

$$y = f(x) = a_{\boldsymbol{s}_m o} - \sum_{i=1}^{a_{\boldsymbol{s}_m o}} \sum_{w=1}^{|\mathcal{H}_k|} \frac{e_{\boldsymbol{h}_w mi} \tilde{a}_{\boldsymbol{h}_w}}{\sum_{l=1}^{|\mathcal{H}_k|} e_{\boldsymbol{h}_l mi} \tilde{a}_{\boldsymbol{h}_l} + e_{\boldsymbol{s}_m mi} x} > \tilde{a}_{\boldsymbol{s}_m}^{(k)}.$$

Furthermore, the derivative

$$
\begin{aligned}
f'(x) &= \sum_{i=1}^{a_{\boldsymbol{s}_m o}} \sum_{w=1}^{|\mathcal{H}_k|} \frac{e_{\boldsymbol{h}_w mi} \tilde{a}_{\boldsymbol{h}_w} e_{\boldsymbol{s}_m mi}}{\left(\sum_{l=1}^{|\mathcal{H}_k|} e_{\boldsymbol{h}_l mi} \tilde{a}_{\boldsymbol{h}_l} + e_{\boldsymbol{s}_m mi} x\right)^2} \\
&= \sum_{i=1}^{a_{\boldsymbol{s}_m o}} \frac{e_{\boldsymbol{s}_m mi}}{\left(\sum_{l=1}^{|\mathcal{H}_k|} e_{\boldsymbol{h}_l mi} \tilde{a}_{\boldsymbol{h}_l} + e_{\boldsymbol{s}_m mi} x\right)} \sum_{w=1}^{|\mathcal{H}_k|} \frac{e_{\boldsymbol{h}_w mi} \tilde{a}_{\boldsymbol{h}_w}}{\left(\sum_{l=1}^{|\mathcal{H}_k|} e_{\boldsymbol{h}_l mi} \tilde{a}_{\boldsymbol{h}_l} + e_{\boldsymbol{s}_m mi} x\right)} \\
&= \sum_{i=1}^{a_{\boldsymbol{s}_m o}} \frac{\Pr\left(\boldsymbol{s}_m \mid \boldsymbol{Z}_{mi} = \boldsymbol{s}_m, \boldsymbol{q}_{mi}; x\right)}{x} \Pr\left(\boldsymbol{s}_m \mid \boldsymbol{Z}_{mi} \in \mathcal{H}_k, \boldsymbol{q}_{mi}; x\right) \\
&< \sum_{i=1}^{a_{\boldsymbol{s}_m o}} \frac{\Pr\left(\boldsymbol{s}_m \mid \boldsymbol{Z}_{mi} = \boldsymbol{s}_m, \boldsymbol{q}_{mi}; x\right)}{x} \\
&< 1.
\end{aligned}
$$

Both inequalities are strict when quality scores are positive and finite because then $e_{\boldsymbol{s}_m mi} < 1$. Thus, fixed point iteration of $f(x)$ from initial point $x_0 > \tilde{a}^{(k)}_{\boldsymbol{s}_m}$ converges monotonically to $\tilde{a}^{(k)}_{\boldsymbol{s}_m}$. Since $a_{\boldsymbol{s}_m o} \geq \tilde{a}^{(k)}_{\boldsymbol{s}_m}$, we can initialize with $x_0 = a_{\boldsymbol{s}_m o}$ to guarantee convergence.

#### S4 Error Estimation

For accurate differentiation of sequencing errors from true variation, it is important to estimate the error rates for every NGS sample [4, 7]. Thus, AmpliCI should be used to estimate error profiles of the current sample before denoising it. Like DADA2 [4], we assume the expected error rate per position is determined by the substitution type and the quality score.

The error estimation process of AmpliCI alternates between estimating haplotypes and estimating errors. First, error rates for each quality score and substitution type are initialized to those dictated by the Phred quality scores [8] of Eq. (6) and assuming equal substitution probabilities, $\gamma_{hr} = \frac{1}{3}$ for all $h, r \in \{\mathrm{A}, \mathrm{C}, \mathrm{G}, \mathrm{T}\}, h \neq r$ and quality score $q$. In the main loop, AmpliCI is iterated once to add another haplotype to set $\mathcal{H}_k$ at iteration $k$. Conditional on $\mathcal{H}_k$, we use the estimated posterior probability Eq. (S5) to assign all reads to one haplotype source and count the occurrences of each error type (quality score, haplotype nucleotide, read nucleotide combination). For each of the source nucleotides $h$, count-weighted LOESS regression is applied to estimate the expected log error rate $\widehat{\Pr}(r \mid h; q)$ for each read base $r \neq h$ as a function of quality score $q$. Following DADA2 [4], we truncate these estimated probabilities at 0.25 above and $10^{-7}$ below. Then, non-error rates are updated as

$$\widehat{\Pr}(h \mid h; q) = 1 - \sum_{r \neq h} \widehat{\Pr}(r \mid h; q).$$

After enough haplotypes have been selected so that at least half of the reads have a maximum conditional log likelihood $\max_{\boldsymbol{h}_l \in \mathcal{H}_k} \pi_l \Pr(\boldsymbol{r}_i \mid \boldsymbol{h}_l; \boldsymbol{q}_i)$ above a user-defined threshold (default: $-100$), subsequent iterations of AmpliCI are performed with the most recent estimate of the error rates. We found this delayed use of estimated error rates necessary to avoid premature termination of the algorithm due to inflation of the estimated error rates when there are still too few haplotypes. We continue to add haplotypes and re-estimate error rates until they stabilize. To assess convergence, for every possible quality score, we compute the cosine distance between occurrence counts of each combination of haplotype nucleotide and read nucleotide before and after addition of a new haplotype, and stop iterations when the minimum cosine distance across quality scores exceeds a threshold (default = 0.999). The final estimated error profile is output to file, and AmpliCI is run again with the fixed error profile given in the file. Typically, the time spent for error estimation is much less than the time spent denoising (see Table 2 in the main text).

#### S5 Bayesian Information Criterion (BIC) for Haplotype Selection

The calculation of BIC is based on a hierarchical mixture model, where we model the dependence between the current haplotype sequences. The collection of $k$ haplotypes $\mathcal{H}_k$ is parameter-rich, but in most applications also highly dependent because of biological relatedness. We penalize the excess parameterization by assuming the haplotypes in $\mathcal{H}_k$ are independent realizations of a Jukes-Cantor [9] model over evolutionary times $\boldsymbol{v} = (v_1, v_2, \ldots, v_k)$ from a shared ancestor $\boldsymbol{a} = (a_1, a_2, \ldots)$, with $j$th nucleotide $a_j$. Thus, the hierarchical likelihood $L(\boldsymbol{a}, \boldsymbol{v}|\mathcal{H}_k)$ for haplotype sequences $\mathcal{H}_k$ is a product of

$$\Pr(\boldsymbol{h}_i; \boldsymbol{a}, v_i) = \prod_{j=1}^{l} \left(\frac{1}{4} + \frac{3}{4}e^{-4v_i/3}\right)^{1\{a_j = h_{ij}\}} \left(\frac{1}{4} - \frac{1}{4}e^{-4v_i/3}\right)^{1\{a_j \neq h_{ij}\}}. \tag{S2}$$

for $i = 1, 2, \ldots, k$. When doing the inference, we directly plug the method-of-moments estimators (MMEs) of $\boldsymbol{v}$ conditional on estimated $\mathcal{H}_k$ into the likelihood function. The nucleotides in the ancestral sequence $\boldsymbol{a}$ are set to the modal nucleotide observed at each site in $\mathcal{H}_k$. We do not account for indels among the haplotypes, as doing so would require an expensive multiple sequence alignment. When indels actually exist among the true haplotype variants, this simplification will lead to overestimation of $v_i$ for indel-containing haplotypes and a chance they will be excluded by BIC.

Suppose $n_p^* = l + 2k - 1 + 12n_q$ is the total number of parameters involved in the hierarchical model, where $l$ is the maximum haplotype length, $k$ is the current number of clusters, and $n_q$ is the range of quality scores observed in the data ($n_q = 0$ if assuming Phred quality scores). Then the BIC for the hierarchical mixture model is

$$\ln(n)n_p^* - 2\arg\max_{\boldsymbol{\theta}} \ln\left[L(\boldsymbol{\theta} \mid \mathcal{R})\right] - 2\arg\max_{\boldsymbol{a},\boldsymbol{v}} \ln\left[L(\boldsymbol{a},\boldsymbol{v}|\mathcal{H}_k)\right]. \tag{S3}$$

The model log likelihood $\ln[L(\boldsymbol{\theta} \mid \mathcal{R})]$ is given by (1) in the main text after substituting in a hard-clustering approximation to $\pi_k$. Specifically, each read $\boldsymbol{r}_i \in \mathcal{R}$ is assumed to originate from the haplotype $\boldsymbol{h}_k$ for which $\Pr(\boldsymbol{r}_i \mid \boldsymbol{Z}_i = \boldsymbol{h}_k; \boldsymbol{q}_i)$ is maximized. If the current source haplotype assigned to read $\boldsymbol{r}_i$ is $\hat{\boldsymbol{z}}_i$, then the MLE for the mixing proportion is estimated as

$$\hat{\pi}_k \approx \frac{1}{n}\sum_{i=1}^{n} 1_{\{\hat{\boldsymbol{z}}_i = \boldsymbol{h}_k\}}.$$

Maximization of the hierarchical log likelihood in Eq. (S3) is approximate, both because the two terms are maximized independently and because they are maximized approximately: $\{\mathcal{H}, \boldsymbol{\theta}_q\}$ via the AmpliCI algorithm and $\boldsymbol{\pi}$, $\boldsymbol{a}$ and $\boldsymbol{v}$ via the approximations described here.

#### S6 Diagnostic for Contamination

Real data are replete with contamination [10]. When a haplotype $\boldsymbol{s}_m$ is selected for inclusion in $\mathcal{H}_k$, we wish to assess the strength of the evidence for its existence as a haplotype rather than a contaminant. If sequence $\boldsymbol{s}_m$ is a real haplotype, then the $a_{\boldsymbol{s}_m o} = A_{\boldsymbol{s}_m} + N_{\mathcal{H}_k \boldsymbol{s}_m}$ observed copies of $\boldsymbol{s}_m$ are the sum of the number $A_{\boldsymbol{s}_m}$ of error-free reads of $\boldsymbol{s}_m$ (whose expectation is the scaled abundance $a_{\boldsymbol{s}_m}$) and the number of misreads $N_{\mathcal{H}_k \boldsymbol{s}_m}$ from true haplotypes in $\mathcal{H}_k$. If reads are conditionally independent, then $N_{\mathcal{H}_k \boldsymbol{s}_m}$, given the observed count $a_{\boldsymbol{s}_m o}$, has a Poisson binomial distribution with probability mass function

$$P\left(N_{\mathcal{H}_k \boldsymbol{s}_m} = y \mid a_{\boldsymbol{s}_m o}, \mathcal{H}_k\right) = \sum_{\sum_i d_{mi} = y} \prod_{i=1}^{a_{\boldsymbol{s}_m o}} p_{mi}^{1\{d_{mi}=1\}} (1 - p_{mi})^{1\{d_{mi}=0\}},$$

where $\boldsymbol{d}_m$ is a vector of $a_{\boldsymbol{s}_m o}$ indicators of the event that $\boldsymbol{r}_{mi}$ is a misread and

$$p_{mi} = \frac{\sum_{w=1}^{|\mathcal{H}_k|} e_{\boldsymbol{h}_w mi} a_{\boldsymbol{h}_w}}{\sum_{w=1}^{|\mathcal{H}_k|} e_{\boldsymbol{h}_w mi} a_{\boldsymbol{h}_w} + e_{\boldsymbol{s}_m mi} a_{\boldsymbol{s}_m}} \tag{S4}$$

is the probability that the $i$th read $\boldsymbol{r}_{mi}$ matching sequence $\boldsymbol{s}_m$ is a misread from current haplotype set $\mathcal{H}_k$.

If sequence $\boldsymbol{s}_m$ includes contaminating reads, then $a_{\boldsymbol{s}_m o} = Z_m + A_{\boldsymbol{s}_m} + N_{\mathcal{H}_k \boldsymbol{s}_m}$, where $Z_m$ is the number of contaminating copies of $\boldsymbol{s}_m$. It is not clear how to model the contamination process $Z_m$, so we assume a very simple process that generates a fixed $Z_m = z$ copies. In particular, we assume $z = c - 1$, where $c$ is the abundance threshold provided by the user. At the default value $c = 2$, we assume 1 copy of sequence $\boldsymbol{s}_m$ is produced by contamination. To assess whether $\boldsymbol{s}_m$ is a contaminant, we compute the diagnostic probability that all but $z$ observed copies of $\boldsymbol{s}_m$ are misreads of the haplotype set $\mathcal{H}_k$,

$$\Pr\left(N_{\mathcal{H}_k \boldsymbol{s}_m} \geq a_{\boldsymbol{s}_m o} - z \mid a_{\boldsymbol{s}_m o}, \mathcal{H}_k\right) = \sum_{i=a_{\boldsymbol{s}_m o}-z}^{a_{\boldsymbol{s}_m o}} \Pr\left(N_{\mathcal{H}_k \boldsymbol{s}_m} = i \mid a_{\boldsymbol{s}_m o}, \mathcal{H}_k\right),$$

where we substitute in the estimates $\tilde{a}_{\boldsymbol{h}_w}$ and $\tilde{a}_{\boldsymbol{s}_m}^{(k)}$ for the unknown abundance values in Eq. (S4). We use a generalization of Pascal's triangle for binomial coefficients to compute the Poisson binomial probabilities in the sum [11]. This calculation becomes expensive as $a_{\boldsymbol{s}_m o}$ increases, so when $a_{\boldsymbol{s}_m o} > 100$, the diagnostic probability is instead computed using the normal approximation [12]. Small values of the diagnostic probability suggest $\boldsymbol{s}_m$ is a real haplotype since it is difficult to explain $a_{\boldsymbol{s}_m o}$ observed reads merely as misreads of haplotypes in $\mathcal{H}_k$, even after allowing for some low-level contamination $z$. To avoid over-engineering AmpliCI, we did not optimize a threshold for this probability. Instead, we chose threshold $0.001/M$, adjusting for the original $M$ candidate haplotypes, unique sequences with observed abundance exceeding $c$. This choice is coded as the default value in AmpliCI and was used in all simulation, mock, and real datasets analyzed. This threshold has proven to be a liberal choice, and AmpliCI-con, where the threshold is set to $1 \times 10^{-40}$ and applied post hoc to the default AmpliCI output, generally produces better results. This conservative threshold is inspired by the significance level used in the DADA2 algorithm when partitioning a cluster [4].

### S7 Abundance Estimation

AmpliCI reports the estimated scaled abundances $\{\tilde{a}_{\boldsymbol{h}_1}, \tilde{a}_{\boldsymbol{h}_2}, \ldots, \tilde{a}_{\boldsymbol{h}_K}\}$ as the haplotype abundance values for downstream analyses. These values underestimate the true abundances by an unknown misread rate. Under the AmpliCI assumption that the misread rate is constant regardless of source haplotype, these abundance values are proportionally correct. Unfortunately, the constant misread rate assumption is probably not valid [13]. Without mock datasets where the true abundances are actually known [14], it can be difficult to assess the effect of this assumption. In this section, we discuss methods to estimate the true abundance values for each detected haplotype, while possibly adjusting for variation in the misread rate. The ideas have not been tested.

A uniform misread rate is hypothetically estimable as $\hat{\alpha} = 1 - \sum_{k=1}^{K} \tilde{a}_{\boldsymbol{h}_k}/n$ for $n$ reads, allowing estimation of true abundances as $\tilde{a}_{\boldsymbol{h}_k}/(1 - \hat{\alpha})$ for each $k$. When the misread rates vary by sequence, AmpliCI abundance estimates may be correctable through a post-hoc adjustment. A true mixture model can partition evidence for the $K$ haplotypes using the posterior assignment probabilities,

$$\Pr\left(\boldsymbol{Z}_i = \boldsymbol{h}_k \mid \boldsymbol{r}_i; \boldsymbol{q}_i\right) = \frac{\pi_k \Pr(\boldsymbol{r}_i \mid \boldsymbol{Z}_i = \boldsymbol{h}_k; \boldsymbol{q}_i)}{\sum_{w=1}^{K} \pi_w \Pr(\boldsymbol{r}_i \mid \boldsymbol{Z}_i = \boldsymbol{h}_w; \boldsymbol{q}_i)}, \text{ for } k = 1, 2, \ldots, K, \tag{S5}$$

where $\boldsymbol{Z}_i$ is the unknown haplotype source of the $i$th read $\boldsymbol{r}_i$. When all parameters are at their maximum likelihood estimates, the maximum likelihood abundance estimate of haplotype $k$ is

$$n\hat{\pi}_k = \sum_{i=1}^{n} \widehat{\Pr}\left(\boldsymbol{Z}_i = \boldsymbol{h}_k \mid \boldsymbol{r}_i; \boldsymbol{q}_i\right). \tag{S6}$$

Further, any read $\boldsymbol{r}_i$ can be assigned to the $\boldsymbol{h}_k$ that maximizes (S5) to obtain a hard-clustering of reads. AmpliCI does not obtain MLEs and hence not equality (S6), but after approximate error parameters have been estimated (see §S4), it can iterate Eqs. (S5) and (S6) starting from $\pi_k \propto \tilde{a}_{\boldsymbol{h}_k}$. Such iteration describes an EM algorithm to estimate $\hat{\boldsymbol{\pi}}$, conditional on error parameters and haplotypes $\mathcal{H}_K$, under a model that makes no assumption of constant misread rate. Indeed, AmpliCI could be used to estimate $K$ and provide initial estimates for an EM algorithm that estimates MLEs of not only mixing proportions $\boldsymbol{\pi}$, but also error parameters and haplotypes $\mathcal{H}_K$.

In real samples, where contamination is common, some reads are contaminants and belong in no haplotype cluster. Having eliminated some candidates using the contamination diagnostic (§S6), we could use the resulting subset of non-contaminant reads in the calculations above. Unfortunately, there remain many other low abundance reads, both true haplotypes and contaminants, which have never been diagnosed for contamination. The best strategy would make a principled decision about whether each read is a contaminant by formulating a more realistic contamination model. We leave further development to future research.

### S8 Deeper Analysis of Mock Datasets

Since real datasets often contain contaminants from untargeted species [5], it is reasonable to compare estimated haplotypes not only to the mock community references *but also* to an outside 16S rRNA gene database (Supplementary Table S2). One way to assess predicted haplotypes when there are true contaminants is to assume true contaminants are unlikely to be similar to mock community species [5]. Under this assumption, haplotypes that are $> 97\%$ similar, but $< 100\%$ similar to mock reference sequences are counted as false positives, because they are *most* likely to be unrecognized misreads of mock references. This approach is not foolproof since such sequences may be authentic, previously unrecognized biological variants of the mock community species. Meanwhile, haplotypes that are $< 97\%$

similar to mock references, but 100% similar to database sequences, are likely true contaminants that should not be counted against the denoiser. Assuming the database is exhaustive, those that are $> 97\%$ similar, but not 100% similar to members of the Silva v132 rRNA gene database [2] would be false positives. However, assessment of haplotypes against any database is problematic, because databases are imperfect and messy [15]. Databases serve as repositories for sequences with sequencing errors, and are incomplete, lacking many of the real biological variants that exist for many bacterial species [16]. Thus, comparison to a database can inflate both true positive and false positive rates in unpredictable ways. Finally, it is completely unclear how to count unmatched sequences with $< 97\%$ similarity to all known reference sequences. Nevertheless, Table S2 shows AmpliCI usually has fewer or no more false positives than other methods under this assessment scheme.

#### S9 The Challenges in Comparing Denoising Algorithms

The pipelines for denoising 16S rRNA gene datasets are quite variable (Supplementary Table S3), and there is no consensus on the best pipeline. Each of these methods, except AmpliCI, is distributed with a complete pipeline and set of use recommendations, making it extremely difficult to accurately compare performance of the denoising component of the pipeline. We have tried to test the methods on mock data within a universal pipeline recommended by DADA2 and Deblur. All denoisers are applied to forward reads after demultiplexing and quality control. After denoising, chimaera sequences are removed by the UCHIME3 *de novo* method [1]. However, there were persistent differences. For chimaera removal, UNOISE3 uses an embedded UCHIME3 *de novo* to detect chimaeras (see § 3.2), and stand-alone UCHIME3 *de novo* may lead to higher error rates (https://drive5.com/usearch/manual/uchime_algo.html), suggesting that the embedded and stand-alone tools may not be the same. Another difference was Deblur's prefiltering step against the Greengenes v13.8 database [17] to remove reads that are unlikely to arise from any 16S rRNA gene. Since we could not turn off this "feature" of Deblur, we ran AmpliCI with and without such filtering.

It is also important to acknowledge that algorithm parameters may be set under the assumption of the tool's recommended pipeline. For example, UNOISE3 is recommended to take merged paired-end reads after stringent quality filtering, while AmpliCI, DADA2, and Deblur all denoise either forward or reverse reads. It seems likely, then, that the error model used by UNOISE3 was trained on merged reads, which are generally higher quality and with error profiles quite different from those observed in forward or reverse reads.

Even if we can isolate the denoising algorithm from its pipeline, it remains difficult to evaluate the performance of the denoising methods. The fact is, we cannot know the true classification of sequences even for the mock data. PCR errors that are heavily amplified will look, in every imaginable way, like real, low level variants. Their only distinguishing feature, under uniform sampling and amplification, is that they should be less abundant than any true variant. The only sure comparison is on simulation data, but they lack realism, especially when it comes to mimicking the level of contamination and error patterns present in real datasets.

#### S10 Technical Limitations for All Denoising Methods

There are technical limitations shared by almost all denoising methods, including AmpliCI. AmpliCI, DADA2 and Deblur officially require reads with the same length, so all these methods recommend truncating reads during preprocessing. It may be worth optimizing the truncation length [18], because it can affect the final results. Additional nucleotides may better resolve haplotypes, but the last few nucleotides contain more errors with unusual patterns [13], which may not be well-modeled by any error model. UNOISE3 supports variable length amplicon sequences, but still highly recommends length trimming, except for fungal ITS amplicons where large natural length variation abounds (https://drive5.com/usearch/manual/pipe_readprep_trim.html). The AmpliCI software can actually accept variable length reads, but the user should consider the reason for the length variation. True indel variation among the haplotypes is well tolerated, but high rates of sequencing indels are not well modeled by the AmpliCI indel error model. Moreover, no denoising algorithm supports ambiguous base calls, but accommodation would be possible in the AmpliCI probability model. All but UNOISE3 recommend against merging paired end reads, but read mergers that update quality scores correctly (*e.g.*, [19] and the UNOISE3 merger) should produce data that works with both AmpliCI and DADA2, which use and learn the meaning of the quality scores.

#### References


[1] R. Edgar, UCHIME2: improved chimera prediction for amplicon sequencing, bioRxiv 74252.

[2] C. Quast, et al., The SILVA and "All-species Living Tree Project (LTP)" taxonomic frameworks, Nucleic Acids Research 42 (2013) D643–D648.

[3] S. F. Altschul, et al., Basic local alignment search tool, Journal of Molecular Biology 215 (1990) 403–410.

[4] B. J. Callahan, et al., DADA2: High-resolution sample inference from Illumina amplicon data., Nature Methods 13 (2016) 581–583.

[5] R. C. Edgar, UNOISE2: improved error-correction for Illumina 16S and ITS amplicon sequencing, bioRxiv 081257.

[6] A. Amir, et al., Deblur rapidly resolves single-nucleotide community sequence patterns, mSystems 2 (2017) e00191–16.

[7] A. McKenna, et al., The Genome Analysis Toolkit: A MapReduce framework for analyzing next-generation DNA sequencing data, Genome Research 20 (2010) 1297–1303.

[8] B. Ewing, P. Green, Base-calling of automated sequencer traces using Phred. II. error probabilities, Genome Research 8 (1998) 186–194.

[9] T. H. Jukes, C. R. Cantor, in: H. N. Munro, J. B. Allison (Eds.), Mammalian Protein Metabolism, Vol. 3, Academic Press, New York, 1969, pp. 21–132.

[10] S. J. Salter, et al., Reagent and laboratory contamination can critically impact sequence-based microbiome analyses, BMC Biology 12 (2014) 87–87.

[11] G. Kallós, A generalization of Pascal's triangle using powers of base numbers, Annales Mathématiques Blaise Pascal 13 (2006) 1–15.

[12] Y. Hong, On computing the distribution function for the Poisson binomial distribution, Computational Statistics and Data Analysis 59 (2013) 41–51.

[13] M. Schirmer, et al., Insight into biases and sequencing errors for amplicon sequencing with the Illumina MiSeq platform, Nucleic Acids Research 43 (2015) e37.

[14] J. M. Bender, et al., Quantification of variation and the impact of biomass in targeted 16S rRNA gene sequencing studies, Microbiome 6 (2018) 155.

[15] R. Edgar, Taxonomy annotation and guide tree errors in 16S rRNA databases, PeerJ 6 (2018) e5030.

[16] J. Ritari, et al., Improved taxonomic assignment of human intestinal 16S rRNA sequences by a dedicated reference database, BMC Genomics 16 (2015) 1056.

[17] T. Z. DeSantis, et al., Greengenes, a chimera-checked 16S rRNA gene database and workbench compatible with ARB, Applied and Environmental Microbiology 72 (2006) 5069–5072.

[18] M. M. Weinstein, et al., FIGARO: An efficient and objective tool for optimizing microbiome rRNA gene trimming parameters, bioRxiv 610394.

[19] G. Renaud, et al., leeHom: adaptor trimming and merging for Illumina sequencing reads., Nucleic Acids Research 42 (2014) e141.
